# Stronger sexual dimorphism in fruit flies may be favored when congeners are present and females actively search for mates

**DOI:** 10.1101/2022.05.17.492314

**Authors:** Alaine C. Hippee, Marc A. Beer, Allen L. Norrbom, Andrew A. Forbes

## Abstract

Why are some species sexually dimorphic while other closely related species are not? When the degree of sexual dimorphism varies within a genus, an integrative phylogenetic approach may help reveal underlying patterns favoring the evolution of dimorphism. While all female flies in genus *Strauzia* – a genus of true fruit flies – share a multiply-banded wing pattern, males of four species have patterns wherein bands have “coalesced” into a continuous dark streak across much of the wing. We find that the origin of coalesced male wing patterns and pronounced differences in male wing shape correlate with the presumed origin of host plant sharing in this genus. A survey of North American Tephritidae finds just three other genera with specialist species that share host plants. Each has one or more congeners with wing patterns unusual for that genus, and just one genus, *Eutreta*, has those unusual wing patterns only in the male sex. *Eutreta* is also the only other genus among this subset wherein, like *Strauzia*, males hold territories while females search for mates. Sharing the same hosts may result in reproductive character displacement, and when coupled with a biology wherein females actively search for males, may specifically favor sexually dimorphic wing patterns.

## Introduction

Sexually dimorphic traits, characters that differ between biological sexes, have long been a focus of biologists fascinated by the problem of how and why selection acts differently on individuals of the same species. Often, dimorphism in one sex results from direct interactions between sexes^1^. Sexual dimorphism can play a role in mating behavior with differences emerging as a result of sexual selection. Specific examples include when dimorphic traits emerge due to sexual signaling mechanisms, including both mate attraction^2-4^ and the evaluation of mate quality^5,6^. In some other cases, sexual dimorphism can be the result of ecological factors unrelated to intersexual interactions, such as when different sexes have different ecological roles, and those roles favor divergent morphologies^1^. Alternatively, the emergence of sexually dimorphic traits can result from interspecific interactions. For instance, reproductive character displacement can occur when congeners are found in close contact, and this is usually ascribed to selection against interspecific hybridization^7^. One pattern resulting from reproductive character displacement is a higher prevalence of sexual dimorphism when species are in sympatry with close relatives than when they are not^8,9^. Discriminating among the many possible hypotheses to explain the evolution of sexual dimorphism in any given species or genus can be challenging because objectively evaluating all potential explanations may often require a complete accounting of the biology, ecology, behavior, and evolutionary history of the focal group. However, when much of this information is known, and when the presence or degree of sexual dimorphism can be measured across a single genus, it is possible to develop an integrative phylogenetic understanding of the evolution of dimorphic traits (e.g., 10; 11).

Flies in the genus *Strauzia* Robineau-Desvoidy (Diptera: Tephritidae) provide a new opportunity to integrate morphology, phylogeny, behavior, and ecology towards understanding the evolution of sexual dimorphism. *Strauzia* have a long history of taxonomic uncertainty, with a considerable degree of apparent intraspecific variation in several putative species^12^. Recent phylogenetic work has clarified that some of this perceived variation is actually interspecific: while the majority of *Strauzia* species are the lone fly from their genus feeding on any given plant host, in each of two instances three *Strauzia* species share the same plant host^13^. Though this work has improved the taxonomy of the genus, it has also confirmed that some traits are sexually dimorphic, and that at least two traits – wing shape and pattern – are strongly dimorphic in some *Strauzia* species but less variable in others.

Like most true fruit flies^12,14,15^, all *Strauzia* have distinctive darkened patterns on their wings, in most species comprised by orange to moderately brown bands. The most common pattern, shared across females and males of most *Strauzia* species, is the “F-pattern”^16^where the bands on the distal third or more of the wing form an “F” (Figure 1a-c). The “F” is conserved in all female *Strauzia*, except for one species (*S. arculata* (Loew)) with a slightly modified pattern with most of the elements of the F present, although some species may also have anterior or posterior connections of the F to other wing bands. Conservation of the F-pattern may be the result of natural selection: similar banding patterns in other Tephritidae mimic the appearance of jumping spiders and offer protection from predation by those same spiders^17-19^. Though this putatively beneficial trait is otherwise strongly conserved across the genus, the males of four *Strauzia* species instead have a “coalesced” wing pattern, wherein the wing is predominantly occupied by a broad dark brown marking running longitudinally down the wing, with the bands that would otherwise constitute the apical “F” fused, sometimes shortened posteriorly and not or at most partially recognizable (Figures 1d, 3). This coalesced patterning is also often a noticeably darker brown than the wing pattern of the conspecific female. Such wing dimorphism is not only unusual in *Strauzia*, but among Tephritidae generally.

**Figure 1.**
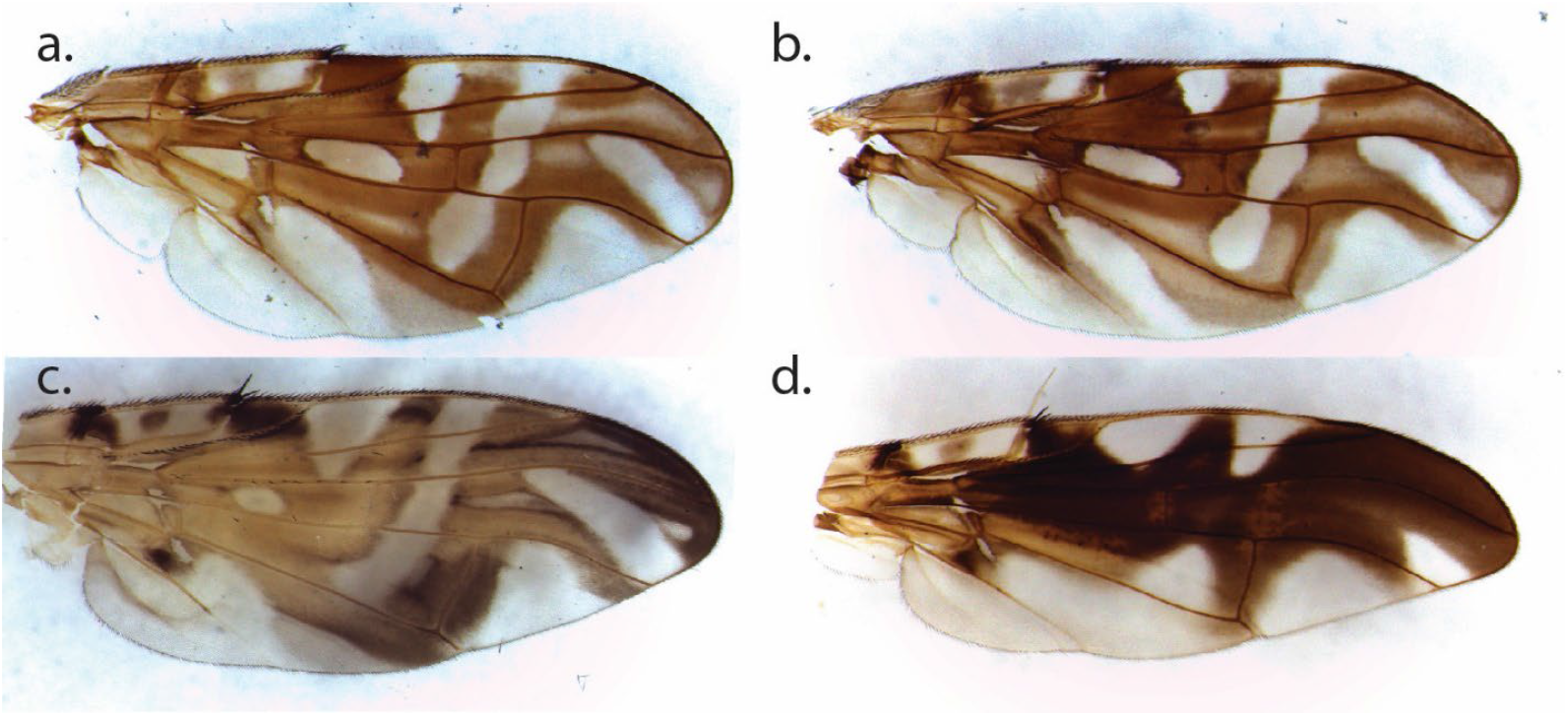
Female and male wings from two representative *Strauzia* species. Wing patterns are generally similar in most species. In *Strauzia intermedia*, for example, females (a) and males (b) both have the typical “F” banding pattern. In other species, like *Strauzia noctipennis*, females (c) have the F pattern, while male wings (d) have a “coalesced” pattern that is darker and more continuous across the center of the wing.

Associations with host plants may also be relevant to *Strauzia* wing evolution. All *Strauzia* species have univoltine life cycles intimately tied to their plant host: males stake out territories on plant leaves, females search among plants to find males, eggs are laid in the apical meristem of the plant, larvae feed on the pith, and pupariation occurs either in the lower stem, root, or soil directly around the plant. In two cases, three species of *Strauzia* specialize on the same host plant. Jerusalem artichoke (*Helianthus tuberosus* L.) is host to *Strauzia longipennis* (Wiedemann), *Strauzia vittigera* (Loew), and *Strauzia longitudinalis* (Loew), while *Strauzia arculata* Steyskal, *Strauzia noctipennis* Stoltzfus, and “Bush’s Fly” (a species not yet formally named and described) all share the sawtooth sunflower (*Helianthus grosseserratus* Martens)^13,20^. As predicted for closely related species that largely overlap in the same habitat^21-24^, previous work has identified evidence of apparent character displacement among the three species of *Strauzia* that share the *H. tuberosus* host, most notably in the form of differences in adult emergence timing^20^. New genetic support for *Strauzia* species limits also demonstrates that three of the four *Strauzia* species with coalesced male wing patterns are among the fly species that share plant hosts with congeners^13,20^.

In this study, we leverage the *Strauzia* phylogeny alongside new morphometric data and previous work detailing their respective host associations, mate choice behaviors, and phenology, to characterize the evolution of sexually dimorphic wing pattern and shape. We also review host association, behavior, and wing dimorphism across other North American Tephritidae to assess whether patterns found in *Strauzia* are representative of a larger theme across the true fruit flies.

## Methods

### Adult Fly Collections and Wing Mounting

From 2011-2021, we collected adult *Strauzia* representing 11 of the 12 named species, plus the undescribed “Bush’s Fly” and another undescribed species that is sister to *S. vittigera* reared from *Helianthus strumosus* (“*strumosus* Fly”). We captured adult flies individually in plastic cups while they rested on host plants. Some flies were also reared from pupae that we had dissected from plant stems and artificially overwintered for 4 months in a refrigerator at 4-8°C. We removed pupae from the refrigerator after 4 months, held them at 18°C for 1 week, and then moved them to a light- and temperature-controlled incubator (16:8 photoperiod; 25°C) to encourage eclosion of adults. All flies were preserved in 95% ethanol and stored at -80°C until use. Only adult flies with wings that were intact or nearly intact were included in the dataset (Supplemental Table 1 – flies included in study).

We removed both fly wings using fine point tweezers and preferentially selected the most intact wing for analysis. We mounted wings on glass slides by soaking each wing in a NaOH solution at 100°C for 1 minute, followed by a 1-minute soak in 95% ethanol - a modified version of the protocol described in Steyskal et al.^25^. Using featherweight tweezers, we gently placed the wing on a glass slide, allowed the remaining ethanol to evaporate, and mounted the wing with several drops of warmed Euparal (BioQuip Products Inc, Rancho Dominguez, California, USA) and a glass coverslip. We allowed the slides to dry on a slide warmer for approximately one week before taking pictures of the slides. Due to changes in product availability, 66 wings were mounted using Permount (Thermo Fisher Scientific, Waltham, MA, USA) instead of Euparal and then were allowed to dry for 48 hours at room temperature prior to wing photography. In total, we analyzed 254 wing slides including 211 slides that we mounted and 43 additional slides that were provided by Dr. Marty Condon (Cornell College, Mt. Vernon, IA).

### Wing Morphometrics and Centroid Analysis

We photographed all *Strauzia* wings using a Leica IC80 HD camera linked to a Leica M125 microscope (Leica Microsystems, Wetzlar, Germany) set to 2X magnification. We opened each wing image in ImageJ v1.52a^26^, then converted images to grayscale, adjusted the orientation so each wing was facing the same direction, and checked to make sure the number of pixels was identical across all images. Then, using the landmark tool in ImageJ, we laid eight single point landmarks that represented the most consistent vein intersections across all *Strauzia* wings (Figure 2). These landmarks were based on a previous set of fourteen landmarks used in analyses of other tephritid fly wings^27^, but we eliminated six landmarks because we failed to find consistent vein intersections across all *Strauzia* species. To avoid variation introduced by different researchers laying the landmarks, one person (ACH) completed all wing landmark analyses. To further eliminate variation introduced by the landmarking process, each wing was landmarked twice, on two separate occasions and in a random order. Then, we compared both sets of landmarks, and the wing sample was eliminated from the analysis if the landmark coordinates differed by more than 1% across the two sets of landmarks for each individual fly. If the sample passed this accuracy threshold, the two sets of landmark coordinates were averaged together to generate a single set of eight coordinates for each wing sample. To test for differences between left and right wings, we mounted both wings from the same male *S. vittigera* (n = 6) and *S. longitudinalis* (n = 5) flies and compared landmarks using the MANOVA statistical procedures described below. We found no difference (P[*vittigera*] = 0.95; P[*longitudinalis*] = 0.99), providing justification for using either wing in subsequent tests, particularly when one wing had been damaged before capture in the wild or during occasional failed slide mounting.

**Figure 2.**
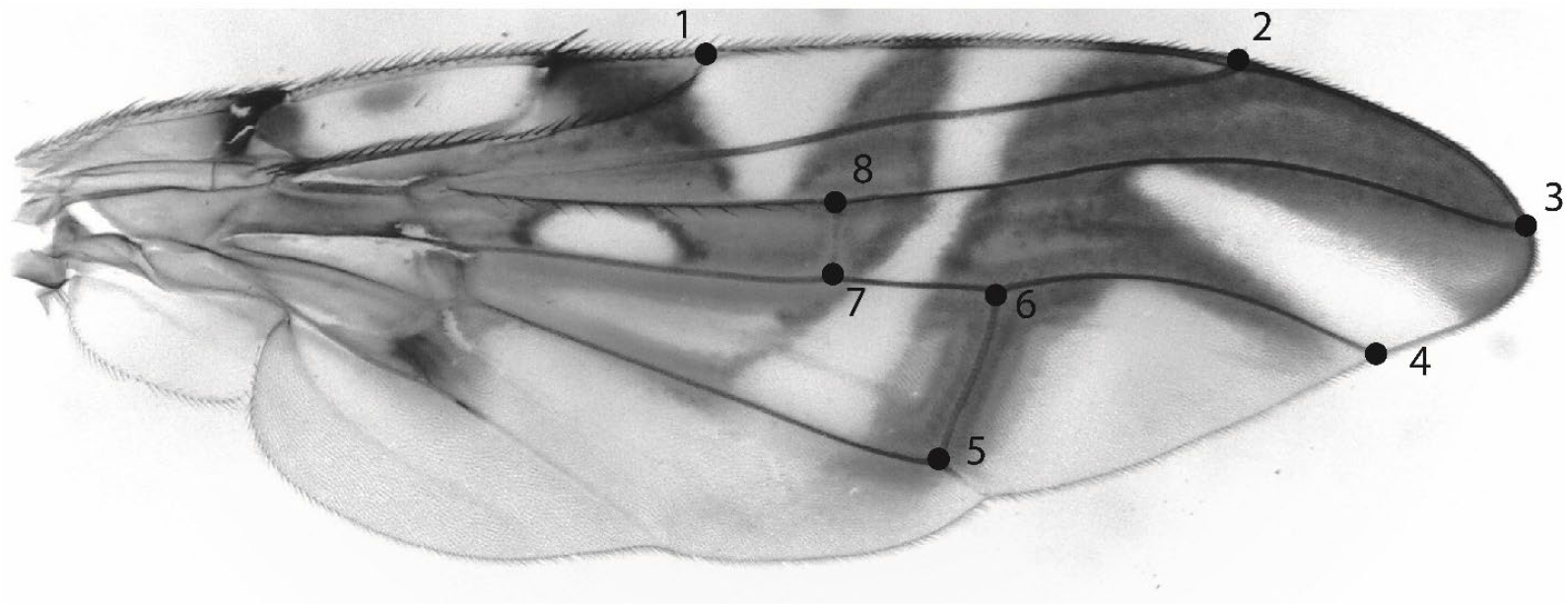
Example *Strauzia* wing with landmarks. Black circles indicate vein intersections used as landmarks for wing morphometric analysis. Numbers next to each circle indicate the landmark number. Landmark locations are based on those described in Marsteller et al^27^.

We imported our landmark coordinates into geomorph v4.0.1^28^ implemented in R to complete a series of wing morphometric analyses. First, we used a generalized Procrustes analysis (GPA^29,30^) using *gpagen* to align the coordinates of all samples using a least squares criterion and projected the resulting coordinates on a linear tangent space^26,31^. Completing the GPA eliminates existing variation due to size, position, and orientation in the landmarks, allowing all remaining variation in landmarks to describe shape differences^31^. The resulting landmarks can be used for multivariate statistical comparisons of shape. We used Principal Component Analyses (PCA) to visualize shape variation between males and females of each *Strauzia* species. All six *Strauzia* species that share plant hosts and four species that do not share hosts were analyzed. For three additional species (*S. rugosum, S. uvedaliae*, and *S. verbesinae*), fewer than three male or female wings were available, which was too few for statistical comparison. We repeated this procedure using the program PAST v4.04^32^ to verify that different morphometrics programs produce similar results.

To determine if wing shape was significantly different between males and females of the same species, we generated a multivariate analysis of variance (MANOVA) of principal components generated during the PCA analysis. We used the broken-stick model^33-35^ on the scree plot generated in PAST v4.04 to select only principal components that account for the majority of the variance for our analyses. In most cases, this was between 2-4 principal components in each analysis. For some comparisons, only one principal component was selected from the broken-stick model. For those cases, a t-test was used to compare the principal components, and we also did a MANOVA by including a second principal component despite it not meeting the broken-stick model criteria. To visualize the magnitude and direction of wing shape change, we used *mshape* in geomorph v4.0.1 to calculate the mean male and female wing shape for each species. Then, using *plotReftoTarget*, we generated points and vectors showing how each wing landmark differs between females and males for each species.

We calculated centroid size for males and females of each species using PAST v4.04 and geomorph v4.0.1. We tested for differences in centroid size between males and females of the same species using t-tests and calculated the average centroid size for males and females of each species. To determine if centroid size differed between *Strauzia* species independently of body size, for all male *S. longipennis, S. longitudinalis, S. perfecta, S. intermedia, S. vittigera, S. noctipennis*, “Bush’s Fly”, and *S. arculata* flies we scaled fly wing centroid sizes by the average fore femur length of each species, as fore femur length has been shown to correlate to body size in Tephritid fruit flies^14^. Indeed, we tested fore femur length and body length size correlation in three species, *S. longipennis, S. longitudinalis*, and *S. vittigera*, and found a strong positive correlation for each species individually (Pearson’s r = 0.89, 0.92, and 0.97 respectively) and combined (Pearson’s r = 0.92). We then tested for a correlation between fore femur length and wing centroid size using log-transformed values for *S. longipennis, S. noctipennis, S. arculata, S. perfecta, S. intermedia, S. vittigera*, and “Bush’s Fly” males.

We also compared wing shape variation among male and female *Strauzia* that share the same host plants. Following the same procedures for principal component analyses and statistical comparison, we compared all males and females that utilize *H. tuberosus* (*S. longipennis, S. vittigera*, and *S. longitudinalis*) and all males and females that share *H. grosseserratus* (*S. arculata, S. noctipennis*, and “Bush’s Fly”) in four separate PCAs and MANOVA procedures.

### Phylogenetic analyses of wing patterns

To contextualize patterns in *Strauzia* wing variation alongside their evolutionary histories, we mapped representative images of male and female wings, PCA plots, and the shape change landmarks on a previously published phylogeny of *Strauzia*^13^. Generated with SNP data from reduced-representation genomics sequencing (3RAD), this phylogeny included 127 *Strauzia* specimens representing 11 of the 12 known *Strauzia* species as well as at least two currently undescribed species.

### Host Sharing and Sexual Dimorphism in other Tephritid Flies

We reviewed the literature pertaining to the biology and morphology of the Tephritidae of the USA and Canada to investigate whether there are common patterns of sexual dimorphism in wing pattern correlated with ecology. We searched the *Handbook of the Fruit Flies (Diptera: Tephritidae) of America North of Mexico*^12^ for all instances where two or more species of the same genus specialized on the same plant host. We narrowly defined a “specialist” fly species as one for which all or most records were from a single plant species. However, we recognize that natural history records might omit geographically restricted or otherwise understudied plant hosts (which would result in flies appearing more specialized than they actually are), or records might include incorrect insect-plant associations, which could result in flies looking more generalist than they actually are. We included instances wherein ≥2 specialist species co-occurred on the same host plant with other, more generalist, congeners but did not include situations where only one specialist species used a plant also used by a congeneric generalist species. Our reasoning in being so restrictive was that we wanted to avoid systems where flies could move to alternative host plants when congeners were locally present.

For each genus identified as having ≥2 specialist species using the same host plant, we then surveyed its respective natural history literature to determine 1) whether any of the host-sharing species showed wing patterns unusual for that genus, 2) whether any species were noted as being sexually dimorphic in wing patterns or shape, and 3) whether both sexes actively searched for mates or if only one sex searched while the other held territories, as is the case in *Strauzia*.

## Results

### Wing Morphometrics and Centroid Analysis

Centroid size measurements showed that wing size is variable among *Strauzia* species and by sex, with *S. intermedia* females having the smallest wings (mean = 874.7 ±70.26), n= 5) and *S. uvedaliae* females having the largest (mean = 1207.2 ±20.32, n = 2). Among comparisons of males and females of the same species, only S. *arculata* (t-test; P-value 0.0005), *S. longitudinalis* (t-test; P-value 0.004), “Bush’s Fly” (t-test; P-value 0.01), and *S. perfecta* (t-test; P-value 0.011) wings had significantly different centroid sizes (Supplemental Table 2). Pairwise comparisons of male and female centroids compared to other species show that the majority (82%) of male and female wing centroids do not differ significantly from each other, with some exceptions in male wings and even fewer among the female comparisons (Supplemental Table 3). Our tests for correlation between fore femur length and centroid size found an overall positive correlation among all species (0.58). Only *S. arculata* had a strong negative correlation between fore femur length and wing centroid size (−0.84). If *S. arculata* is excluded from the pooled analysis, the remaining species have a correlation of 0.62. After scaling male wing centroid size by fore femur length as a proxy for body size, we found even fewer significant differences in centroid size among all possible comparisons across the genus when fore femur measurements were available (Supplemental Table 4). The remaining comparisons that were significantly different all included comparisons with *S. noctipennis* males, which appear to have a significantly different centroid size than *S. arculata*, “Bush’s Fly”, *S. intermedia*, S. *perfecta*, and *S. longipennis*.

Principal Component Analyses (PCAs) of wing shape (with size variation excluded from the comparison) between males and females of the same species showed variation in the presence of wing shape dimorphism across *Strauzia*. Using MANOVAs, we statistically compared the differences between the principal components for each species to determine if wing shape differed significantly among males and females of the same species and among males or females of different species. Three species - *S. intermedia, S. gigantei*, and *S. arculata* - had male and female wings that were not significantly different in shape (Figure 3; Table 1). The remaining *Strauzia* species for which male and female comparisons were possible – *S. noctipennis*, “Bush’s Fly”, *S. longipennis, S. longitudinalis, S. perfecta*, “*strumosus* Fly”, and *S. vittigera* had wing shapes that were significantly different between sexes (Figure 3; Table 1). We also generated vectors showing the direction and magnitude of shape change between the mean female and mean male wing shape of each *Strauzia* species. Across all *Strauzia* wings, male wings were generally narrower and longer than female wings, with *S. noctipennis*, “Bush’s Fly”, *S. longipennis, S. longitudinalis*, “*strumosus* Fly”, and *S. vittigera* showing extreme examples manifested in changes in wing landmarks 1-5 (Figure 3, Supplemental Figure 1).

**Table 1.** MANOVA comparisons of male and female wing shape. The Wilks’ lambda statistic, F value, and P-value are listed for each comparison. The number of principal components included in the analysis based on the results from the broken stick model and the sample size (N) are also included. Bolded rows indicate male and female wings that were significantly different in shape after a correction for multiple comparisons.

| Species | Wilks' lambda | F | P-value | # of PCs included | Males (N) | Females (N) |
| --- | --- | --- | --- | --- | --- | --- |
| *S. intermedia* | 0.5248 | 2.49 | 0.1042 | 4 | 11 | 5 |
| *S. gigantei* | 0.4364 | 3.875 | 0.0831 | 2 | 4 | 5 |
| ***S. vittigera*** | **0.1781** | **30.77** | **>0.0001** | **3** | **27** | **25** |
| **"strumosus Fly"** | **0.1135** | **35.15** | **>0.0001** | **2** | **35** | **18** |
| *S. arcuata* | 0.5604 | 3.922 | 0.0299 | 3 | 15 | 4 |
| ***S. longitudinalis*** | **0.1348** | **157.3** | **>0.0001** | **2** | **9** | **15** |
| **"Bush's Fly"** | **0.1626** | **43.76** | **>0.0001** | **2** | **8** | **4** |
| ***S. noctipennis*** | **0.02645** | **202.4** | **>0.0001** | **2** | **16** | **4** |
| ***S. longipennis*** | **0.1377** | **156.6** | **>0.0001** | **2** | **8** | **6** |
| ***S. perfecta*** | **0.2697** | **14.89** | **0.0007** | **2** | **9** | **5** |

**Figure 3.**
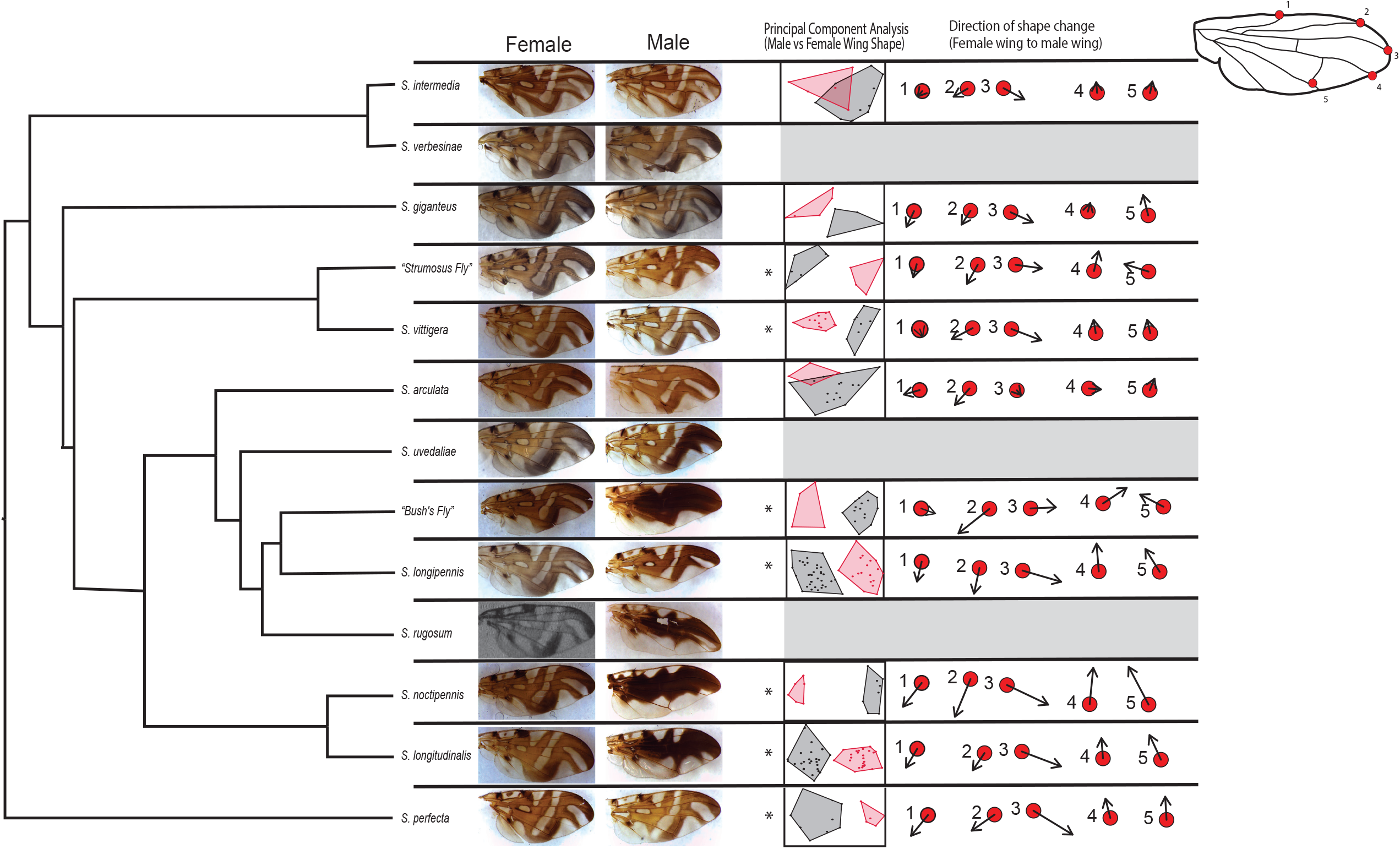
Phylogeny of *Strauzia* with male and female wings. Maximum likelihood phylogeny of *Strauzia* from Hippee et al.^13^ with female (left column) and male (right column) wings next to each *Strauzia* species and the results of the PCA analyses of males (black) and females (red) for each species. Points and arrows on the far right represent the direction and magnitude of shape change between an average female wing (red point) and an average male wing (black arrow). A diagram on the top right shows the location of landmarks 1-5 included in the figure on a standard fly wing. An asterisk next to a PCA plot indicate significantly different male vs. female wing shapes (Table 1). Species with grey shading that are lacking PCA plots were those that did not have adequate sample sizes for analysis. The *S. rugosum* female wing picture is from Stoltzfus 1989.

MANOVAs of wing shape of all *Strauzia* males showed that the majority of *Strauzia* species have significantly different male wing shapes, with *S. noctipennis* and “Bush’s Fly” males differing significantly from those of all other *Strauzia* species included in the analysis (Supplemental Table 6, Supplemental Figure 2). *Strauzia intermedia* males were also significantly different in nine of the eleven species comparisons (Supplemental Table 6). In the *Strauzia* female analysis, the majority (82%) of comparisons did not show a significant difference in shape with “Bush’s Fly” females not having any significant shape differences with the ten other species included in the analysis (Supplemental Table 7; Supplemental Figure 3). *Strauzia longitudinalis* females were the most different, with their shape differing significantly from five of the ten species included in the analysis (Supplemental Table 7).

We also compared the wing shapes of males and females of *Strauzia* species that share the same host plant species in another set of MANOVAs. Three *Strauzia* species – *S. longipennis, S. longitudinalis*, and *S. vittigera* – share the same host plant, *H. tuberosus*. Three additional *Strauzia* species – *S. arculata, S. noctipennis*, and “Bush’s Fly” – also share their host plant, *H. grosseserratus*. All males on *H. tuberosus* and *H. grosseserratus* were significantly different from each other (Supplemental Table 5; Supplemental Figures 4,5). Among the females on *H. tuberosus*, only *S. longipennis* and *S. longitudinalis* were significantly different from each other (P-value 0.003) (Supplemental Table 5; Supplemental Figure 6). On *H. grossererratus, S. arculata* was significantly different from both “*Bush’s Fly*” (P-value 0.01) and *S. noctipennis* (P-value 0.004), but “*Bush’s Fly*” and *S. noctipennis* were not significantly different from each other (P-value >0.05) (Supplemental Table 5, Supplemental Figure 7). Wing shape variation was primarily in landmarks 1 through 5, with landmarks 6 through 8 showing little to no variation between males and females of any *Strauzia* species.

### Phylogeny of Wing Pattern and Shape

We summarized the *Strauzia* phylogeny^13^ into a species tree representing all valid *Strauzia* species, with the exception of *S. stoltzfusi* Steyskal whose host is unknown, plus “*Bush’s Fly*” and the currently undescribed “*strumosus* Fly” (included in the Hippee et al. phylogeny as *S. vittigera* from host *H. strumosus*) (Figure 3). Female wings showed no major pattern variation across the phylogeny with the exception of those of *S. arculata* and “Bush’s Fly”, which both have anterior and posterior connections between the “F” and the more basal wing markings and a near to total loss of connection between the anterior and posterior parts of the “F”, such that it instead takes the form of two chevron shapes at the distal end of the wing (Figure 3).

The male wings of many species show more extreme variation in wing pattern across the genus (Figure 3). Males of “Bush’s Fly”, *S. noctipennis, S. rugosum*, and *S. longitudinalis* all have a fully coalesced wing pattern. Males of two other species, *S. uvedaliae* and *S. longipennis*, also had noticeably darker banding patterns on the apical part of the wing than their respective conspecific females, but their basal bands were still distinct and not coalesced into a single broad marking. All six species with some obvious difference between male and female wing patterns were in the same clade, joined by only *S. arculata* as the exception in having no apparent wing pattern dimorphism. The remaining *Strauzia* species (*S. perfecta, S. intermedia, S. verbesinae, S. gigantei, S. vittigera*, and the undescribed “*strumosus* Fly” collected from *H. strumosus*) lacked obvious sexual dimorphism in wing pattern.

Wing shape differences between male and female flies are also common, with male wings across all species relatively elongate compared with female wings. However, among measured species that do not share hosts with other *Strauzia*, only *S. perfecta* and the “*strumosus* fly” showed significant male-female wing shape differences. Five of six host-sharing *Strauzia* species showed significant male-female wing shape differences and had no overlap in PCA space between male and female wings. Only *S. arculata* co-occurs on a plant with other *Strauzia* species and had overlap in PCA space between males and females (Table 1, Figure 3).

### Host Sharing and Sexual Dimorphism in other Tephritid Flies

In our review of the US and Canadian tephritid genera, we found only three additional genera (*Aciurina, Eutreta*, and *Valentibulla*) with two or more specialists listed as sharing the same host plant. *Aciurina* and *Valentibulla* are closely related genera^12,36^ and thus could be treated as one clade for comparison with *Strauzia*. In *Aciurina*, as in *Strauzia*, two sets of congeners shared two different host plants, such that we identified a total of six cases of a plant species with multiple specialist congeneric tephritid fly associates (Table 2). In all six cases, one or more of the specialist fly species had wing patterns described by other authors as being unusual for that fly genus, either in both sexes, or only in the males (and see below).

**Table 2.** Table summarizing wing pattern and mate finding behavior traits for congeneric Tephritidae that share a single plant host, with specialists (S) and generalists (G) indicated next to species names. *Aciurina* species marked with * are putative specialists – they have occasionally been noted as having other hosts, but these records are unconfirmed, and we consider them questionable. Species with uncertain host ranges do not affect overall trends.

| Genus | Host plant | Relevant species<br>S = specialist<br>G = generalist | Species with<br>particularly<br>divergent<br>wing patterns | Is the divergent<br>wing pattern<br>found in only<br>one sex? | Notes on wings | Biology/Life history |
| --- | --- | --- | --- | --- | --- | --- |
| *Aciurina* | *Ericameria*<br>*nauseosa*<br>(Rubber<br>rabbitbrush) Ψ | *A. bigeloviae* (S)*<br>*A. maculata* (S)*<br>*A. notata* (S)<br>*A. opaca* (S)<br>*A. trilitura* (S)<br>*A. trixa* (S)* | *A. notata*<br>*A. bigeloviae* | No | All six species can be distinguished from one another based on wing pattern, with *A. notata* wings being most different from others (hyaline with a few thin dark patches along veins). *Aciurina bigeloviae* wings have a high degree of intraspecific variation<sup>58</sup> | Males and females both walk on stems and leaves of host plant. When males see a female, they pursue<sup>39</sup>. |
| *Aciurina* | *Chrysothamnus*<br>*viscidiflorus*<br>(Yellow<br>rabbitbrush) | *A. ferruginea* (S)*<br>*A. idahoensis* (S)<br>*A. lutea* (S)*<br>*A. michaeli* (S)<br>*A. semilucida* (S) | *A. idahoensis*<br>*A. semilucida* | No, but see notes<br>next column | Both *A. idahoensis* and *A. semilucida* are unusual among *Aciurina* in having most of the wing surface hyaline with a few dark transverse bands<sup>40,41</sup>. Though all five species on this host have some sexual dimorphism in wing patterns, both sexes of *A. idahoensis* and *A. semilucida* have wings that differ from the more common *Aciurina* wing pattern.§ | Males and females both walk on stems and leaves of host plant. When males see a female, they pursue<sup>39</sup>. |
| *Eutreta* | *Artemisia*<br>*tridentata*<br>(big sagebrush) | *E. divisa* (S)<br>*E. oregona* (S)<br>*E. diana* (G) | *E. divisa*<br>(males) | Yes | While most male and female *Eutreta* have dark wings with small to tiny hyaline spots, male *E. divisa* wings have two diagonal hyaline stripes not seen on the wings of any other *Eutreta* flies, including conspecific females. | Males hold territories and "fight" with wing displays. Females enter territories for mating<sup>37</sup>. |
| *Strauzia* | *Helianthus tuberosus* (Jerusalem artichoke) | *S. longipennis* (G)<br>*S. longitudinalis* (S)<br>*S. vittigera* (S) | *S. longitudinalis* (males) | Yes | Male *S. longitudinalis* wings have “coalesced” bands of dark color across the length of the wing (13; Figure 3 this paper). | Males hold territories on leaves. “*Wing and body movements may be involved in attracting the female.*”<sup>44</sup> |
| *Strauzia* | *Helianthus grosseserratus* (sawtooth sunflower) | *S. arcuata* (S)<br>“*Bush's Fly*” (S)<br>*S. noctipennis* (S) | “*Bush's Fly*” (males)<br>*S. noctipennis* (males) | Yes | Wings of male of “*Bush's Fly*” and *S. noctipennis* both have “coalesced” wing patterns (13; Figure 3 this paper). | Males hold territories on leaves. “*Wing and body movements may be involved in attracting the female.*”<sup>44</sup> |
| *Valentibulla* | *Ericameria nauseosa* (Rubber rabbitbrush) Ψ | *V. californica* (S)<br>*V. dodsoni* (S)<br>*V. steyskali* (S) | *V. dodsoni* | No | *Valentibulla dodsoni* wings are “*the most distinctive in the genus*”<sup>12</sup>, with the more usual hyaline spots coalesced into a large hyaline patch across much of the posterobasal quadrant. | Males and females both walk on stems and leaves of host plant looking for mates<sup>38</sup>. |

Ψ We note that the taxonomy of *Ericameria nauseosa* is uncertain, and that it may be a complex of different host plants rather than a single species, which could mean that some of these congeneric fly species do not directly interact on the same plant.
§ There appears to be some geographic variation in the degree of sexual dimorphism in these flies, though wings of both sexes generally differ from the “usual” *Aciurina* wing. Differences between male and female wing patterns in both *A. idahoensis* and *A. semilucida* are more pronounced in California populations than in Idaho populations<sup>40,41</sup>. Only *S. semilucida* females from California approach the groundplan for genus *Aciurina*<sup>40</sup>.

Across these four genera, flies differed in their reported mate-finding behavior and in whether or not divergent / unusual wing patters occurred in one or both sexes. In *Eutreta*, as in *Strauzia*, males stake out territories on leaves while females fly about the plant in search of males^37^. And as in *Strauzia, Eutreta* that co-occurred on the same plants alongside congeners had a species (*Eutreta divisa*) with sexually dimorphic wings, and with males being the sex with the unusual wing patterns^12^ – males, and not females, have two diagonal hyaline stripes interrupting the otherwise primarily dark wing (Table 2). In *Aciurina* and *Valentibulla*, both male and female flies are described as walking along stems and leaves, with mating occurring when they encounter one another^38,39^. Some host-sharing congeners in these two genera also had wings that differed from each genus’ wing groundplan, but these divergent patterns occurred in both sexes, even though sexual dimorphism within the divergent pattern was sometimes evident^40,41^.

## Discussion

Our collective results refine our understanding of the evolution of wing shape and pattern in *Strauzia* flies specifically and fruit flies generally. They underscore the importance of species interactions in morphological evolution, but also how differences in mating behavior may change the selective landscape and result in different outcomes for males versus females. We discuss our findings first in the context of *Strauzia* alone and then approach a synthesis by incorporating our review of other tephritids.

Sexual dimorphism in wing patterns are exclusive to, and wing shape differences are most pronounced in, the *Strauzia* clade that includes species that share host plants. Three of the four species with strongly coalesced male wing patterns occur on either *H. tuberosus* (*S. longitudinalis*) or *H. grosseserratus* (*S. noctipennis*, “Bush’s Fly”) alongside other specialist *Strauzia*. In addition, one of the two other species with a darkening of the apical “F” on the male wing is also found on *H. tuberosus* (*S. longipennis*). Similarly, five of six species that co-occur with other *Strauzia* have significantly different male vs. female wing shapes (Table 1, Figure 3) and all co-occurring males have wing shapes that differ from each other (Supplemental Table 5).

We submit that the most parsimonious explanation for the evolution of sexually dimorphic wing patterns and shapes in *Strauzia* is reproductive character displacement in the context of shared plant hosts. Two specific causes might drive this character displacement: a) avoidance of combat between interspecific males and/or b) avoidance of costly interspecific mating attempts. In many tephritids, males congregate at lekking sites (here, leaves) and engage in wing waving and head butting, resulting in one of the interacting males being driven away^42^. Male *Strauzia* have been observed to engage in these male-male battles and have elongated setae on their heads that may be related to male-male aggression^38,43^. Wing markings may convey visual signals to rivals, and interspecific differences may help flies avoid conspecific battles.

Alternatively, because female *Strauzia* search for territorial males waiting on plant leaves^44^, female choice is an important component of mating success and could drive character displacement. It can be costly to attempt mating (or worse, hybridize) with a different species due to time and energy wasted^45-47^, the risk of physical damage or mortality during mating^48,49^, or the wasting of gametes to generate unfit hybrids^50-52^. Though pheromone signals are important for finding mates in many tephritids^42^, wing markings and wing movements are known to be important at close range^42,53^, and visual signals are the primary long-range attractant in some genera^54^. Our data alone do not favor one hypothesis over the other for *Strauzia* (though see discussion regarding the broader patterns in the North American Tephritidae below).

*Strauzia rugosum* (which has a fully coalesced male wing pattern) and *S. uvedaliae* (which has a darkened apical “F” in male its wing pattern) both have modified male wings but do not share hosts with other *Strauzia*, apparently belying the idea that wing pattern dimorphism is a result of reproductive character displacement. However, the *Strauzia* phylogeny makes clear that sexually dimorphic wing patterns are not phylogenetically independent (Figure 3). The evolution of differences in wing pattern appears to have occurred either once, with one subsequent loss in the branch leading to *S. arculata*, or twice, with separate origins in the respective ancestors of the *S. noctipennis*/*S. longitudinalis* and the *S. uvedaliae*/”*Bush’s Fly*”/*S. longipennis*/*S. rugosum* clades. Whichever the case, *S. rugosum* and *S. uvedaliae* are embedded in a clade for which the common ancestor probably had a sexually dimorphic wing pattern. Thus, these species are not exceptions to a rule, but likely represent lineages that evolved via shifts to new host plants after wing pattern dimorphism had already evolved.

While reproductive character displacement seems to best fit patterns in *Strauzia*, the lack of phylogenetic independence among species and the potential for morphological variation to be influenced by multiple evolutionary forces, such as sexual selection^55,56^ and genetic drift^57^ over the course of evolutionary history, allows for alternative explanations for the correlation between wing patterns and host sharing. However, when considered alongside our survey of wing patterning and mating behavior for the North American Tephritidae, the reproductive character displacement hypothesis is hard to replace with another. First, the rarity of host sharing among specialist tephritids – we find this in only three other genera (twice in genus *Aciurina*, just as in *Strauzia*) – suggests that use of the same hosts may be generally disfavored. Second, in all six cases where two or more specialist congeners do share the same host plant, at least one species on the host plant has wing pattern differences in one or both sexes that is unusual for the genus (Table 2). For instance, Foote et al.^12^ describe *Valentibulla dodsoni* as having wings that are “the most distinctive in the genus”, while in Steyskal’s^58^ description of *Aciurina idahoensis* he notes the “…very characteristic pattern of the wing…readily distinguishes this species from any other.” Host sharing being consistently correlated with wing morphologies that diverge from a presumed original state supports a general hypothesis for reproductive character displacement driving changes in tephritid wing patterns.

Further, mate-finding behaviors appear to correlate with the expression of divergent wing morphologies in one versus both sexes. One of the non-*Strauzia* genera in Table 2 – *Eutreta* – also includes a species with sexually dimorphic wing patterns. While females of *Eutreta divisa* have wings much like other flies in the genus, male *E. divisa* flies have two diagonal white stripes not seen on the wings of any female *Eutreta* species^12^. *Eutreta* also share a behavioral similarity with *Strauzia*: males hold territories on leaves while females fly to search for mates^59^. In *Valentibulla* and *Aciurina*, by contrast, both males and females walk around the surface of the host plants searching for mates^60^, and when host sharing flies in these genera have divergent wing patterns, those patterns are seen in both sexes (though some sexual dimorphism may still be present). Selection against mating with other congeneric species may therefore favor more extreme wing pattern changes in both sexes when males and females both search for mates, while the same dramatic wing pattern changes may occur just in males when only the female sex is actively searching. These broader patterns of wing pattern evolution in the Tephritidae also place more weight on the idea that female choice, not male aggression, drives dimorphic patterns in *Strauzia*, as there are no records of male-male (or, importantly, female-female) aggressive behaviors in *Valentibulla* or *Aciurina*). One other species in this genus, *Eutreta fenestrata*, also shows striking male-female wing pattern dimorphism (female as *E. modocorum* and male as *Metatephritis fenestrata*)^12^, but we do not know if it shares hosts with congeners (or if its ancestors did); its only reported host, *Artemisia nova* A. Nelson, is shared with the non-specialist *Eutreta diana*.

Some sexually dimorphic specialist tephritids do not share hosts with congeners. Males and females of two *Acidogona* species (*Acidogona dichromata* (Snow) and *Acidogona stecki* Norrbom) are sexually dimorphic, with the *A. dichromata* sexes sufficiently different in their wing patterns to have been described as separate species^15,60^. Again, it is the males of these two *Acidogona* species whose wings differ from the groundplan of the genus. In these cases, dimorphism could be due to factors unrelated to reproductive character displacement. On the other hand, *A. dichromata, A. stecki* and their congener, *Acidogona melanura*, all use hawkweeds (*Hieracium*) as host plants^12,15^. The contemporary distributions of these three species are allopatric^12,15^, such that these flies do not interact in nature, but dimorphism may have evolved during a period of host sharing by these species ancestors, as may have been the case for *Strauzia rugosum* and *S. uvedaliae* (Figure 3). A general conclusion is that to better address these important evolutionary questions requires further study of species limits and biogeographical histories of both tephritid flies and their plant hosts.

A formal test of the specific hypothesis that morphological character displacement evolves when congeners share hosts requires directly measuring selection – possibly via experimental manipulation of fly wings (e.g., 14, 61). There may also be a role for other forms of reproductive isolation to influence wing pattern differences in concert with reproductive character displacement. Previous work on *H. tuberosus*-associated *Strauzia* found evidence of temporal isolation^20^, whereas the three *H. grosseserratus*-associated *Strauzia* flies appear to have greater overlap in phenology (ACH, personal observation). More temporal overlap might select more strongly for character displacement, such as wing pattern differences, that reinforce species boundaries. Future work should also consider these and other reproductive barriers that may be present when multiple species are sharing the same host.

Finally, a chicken or egg problem: do major changes in fruit fly wing patterns result from host sharing, or do they facilitate shifts to already-occupied hosts? Our results suggest both may be true. While we argue that our findings strongly suggest reproductive character displacement as a driver of changes in wing pattern for *Strauzia* and other true fruit flies, morphological differences could represent exaptations, facilitating host sharing after morphological differences had already evolved. In both *Aciurina* and *Strauzia*, two different plant species are host to more than one congener (Table 2). The scarcity of host sharing by specialists in most tephritid genera, juxtaposed against it occurring twice in both of these morphologically diverse genera, suggest that once new wing morphologies evolve, sharing hosts with congeners may pose fewer problems and it may be easier for flies to share plants.

## Acknowledgements

We would like to thank Marty Condon and Kara Middleton for providing some of the wing slides included in this study. We would also like to thank Gunther Hansen for his contributions to the wing slide mounting, Michael Lopez for working on initial wing comparisons, and Robin K. Bagley for discussions regarding the morphometrics methodology. Quin Baine, Ellen Martinson, and Vince Martinson provided indispensable information about the current state of knowledge regarding *Aciurina* flies and their host plants in the Southwestern U.S.A. Finally, we would like to acknowledge the following people for help in collecting the insects used in this manuscript including: Jarod Armenta, Darius Ballard, Elana Becker, Charles Bray at F.W. Kent State Park, Neisha Croffitt, Sarah Duhon, Maren Elnes, Demaceo Howard, Patrick Kelly, Dacia Lipkea, Tom Powell, Emily Reasoner, Halee Schomberg, Bob Smith, Heather Widmayer, Kyle Woods, and Alex Young. Mention of trade names or commercial products in this publication is solely for the purpose of providing specific information and does not imply recommendation or endorsement by the USDA. USDA is an equal opportunity provider and employer.

