## Supplemental Tables and Figures for "Stronger sexual dimorphism in fruit flies may be favored when congeners are present and females actively search for mates"

Supplemental Table 1. A table including collection, species, and morphological information for each fly wing used in these analyses. Columns include sample ID/wing ID, species, sex, collection location (state in USA), and collection site GPS coordinates.

| Wing ID/Fly ID | Species | Sex | Host Plant | Collection Location | Collection Site Latitude | Collection Site Longitude |
|---|---|---|---|---|---|---|
| AF100 | *S. longitudinalis* | Male | *H. tuberosus* | Iowa | 41.650823 | -91.560224 |
| AF101 | *S. longipennis* | Male | *H. tuberosus* | Iowa | 41.650823 | -91.560224 |
| AF103 | *S. longipennis* | Male | *H. tuberosus* | Iowa | 41.650823 | -91.560224 |
| AF104 | *S. longitudinalis* | Male | *H. tuberosus* | Iowa | 41.650823 | -91.560224 |
| AF106 | *S. longipennis* | Male | *H. tuberosus* | Iowa | 41.650823 | -91.560224 |
| AF32 | *S. longipennis* | Male | *H. tuberosus* | Iowa | 41.650823 | -91.560224 |
| AF37 | *S. longipennis* | Male | *H. tuberosus* | Iowa | 41.650823 | -91.560224 |
| AF73 | *S. vittigera* | Male | *H. tuberosus* | Iowa | 41.650823 | -91.560224 |
| AF88 | *S. vittigera* | Male | *H. tuberosus* | Iowa | 41.650823 | -91.560224 |
| AF92 | *S. longitudinalis* | Male | *H. tuberosus* | Iowa | 41.650823 | -91.560224 |
| AF97 | *S. longitudinalis* | Male | *H. tuberosus* | Iowa | 41.650823 | -91.560224 |
| AH1061 | *S. intermedia* | Female | *R. laciniata* | Iowa | 41.747012 | -90.515424 |
| AH1063 | *S. uvedaliae* | Male | *S. uvedalia* | North Carolina | 35.92429 | -79.050225 |
| AH1065 | *S. verbesinae* | Male | *V. occidentalis* | North Carolina | 35.97307 | -78.96777 |
| AH1067 | *S. verbesinae* | Male | *V. occidentalis* | North Carolina | 35.97307 | -78.96777 |
| AH1068 | *S. uvedaliae* | Female | *S. uvedalia* | Virginia | 37.533081 | -77.479227 |
| AH1110 | *S. noctipennis* | Female | *H. grosseserratus* | Iowa | 41.735516 | -91.722503 |
| AH1115 | *S. noctipennis* | Male | *H. grosseserratus* | Iowa | 41.735516 | -91.722503 |
| AH120 | *"Bush's Fly"* | Male | *H. grosseserratus* | Iowa | 43.38149 | -95.1838 |
| AH124 | *"Bush's Fly"* | Male | *H. grosseserratus* | Iowa | 43.38149 | -95.1838 |
| AH125 | *S. arculata* | Male | *H. grosseserratus* | Iowa | 43.38149 | -95.1838 |
| AH132 | *S. arculata* | Male | *H. grosseserratus* | Iowa | 43.38149 | -95.1838 |
| AH136B | *S. arculata* | Male | *H. grosseserratus* | Iowa | 43.38149 | -95.1838 |
| AH142B | *S. arculata* | Male | *H. grosseserratus* | Iowa | 43.38149 | -95.1838 |
| AH144B | *S. arculata* | Male | *H. grosseserratus* | Iowa | 43.38149 | -95.1838 |
| AH146 | *S. arculata* | Male | *H. grosseserratus* | Iowa | 43.38149 | -95.1838 |
| AH147B | *S. arculata* | Male | *H. grosseserratus* | Iowa | 43.38149 | -95.1838 |
| AH149 | *S. arculata* | Male | *H. grosseserratus* | Iowa | 43.38149 | -95.1838 |
| AH156 | *"Bush's Fly"* | Male | *H. grosseserratus* | Iowa | 43.38149 | -95.1838 |
| AH157B | *S. arculata* | Male | *H. grosseserratus* | Iowa | 43.38149 | -95.1838 |
| AH160 | *S. noctipennis* | Female | *H. grosseserratus* | Iowa | 43.38149 | -95.1838 |
| AH168 | *"Bush's Fly"* | Male | *H. grosseserratus* | Iowa | 43.38149 | -95.1838 |
| AH172 | *"Bush's Fly"* | Female | *H. grosseserratus* | Iowa | 43.38149 | -95.1838 |
| AH175 | *"Bush's Fly"* | Male | *H. grosseserratus* | Iowa | 43.38149 | -95.1838 |
| AH177 | *S. arculata* | Male | *H. grosseserratus* | Iowa | 43.38149 | -95.1838 |
| AH182 | *S. arculata* | Male | *H. grosseserratus* | Iowa | 43.38149 | -95.1838 |
| AH183 | *S. arculata* | Female | *H. grosseserratus* | Iowa | 43.38149 | -95.1838 |
| AH186 | *"Bush's Fly"* | Male | *H. grosseserratus* | Iowa | 43.38149 | -95.1838 |
| AH189 | *"Bush's Fly"* | Male | *H. grosseserratus* | Iowa | 43.38149 | -95.1838 |
| AH19 | *"Bush's Fly"* | Male | *H. grosseserratus* | Iowa | 43.38149 | -95.1838 |
| AH193 | *"Bush's Fly"* | Male | *H. grosseserratus* | Iowa | 43.38149 | -95.1838 |
| AH194 | *"Bush's Fly"* | Male | *H. grosseserratus* | Iowa | 43.38149 | -95.1838 |
| AH259 | *S. vittigera* | Male | *H. tuberosus* | Iowa | 41.665195 | -91.513398 |
| AH261 | *S. vittigera* | Male | *H. tuberosus* | Iowa | 41.65288 | -91.480382 |
| AH27 | *S. arculata* | Female | *H. grosseserratus* | Iowa | 43.38149 | -95.1838 |
| AH302 | *S. perfecta* | Male | *A. trifida* | Iowa | 41.695322 | -92.232 |
| AH324 | *S. longitudinalis* | Male | *H. tuberosus* | Iowa | 41.65288 | -91.480382 |
| AH336 | *S. longitudinalis* | Male | *H. tuberosus* | Iowa | 41.65288 | -91.480382 |
| AH339 | *S. longitudinalis* | Male | *H. tuberosus* | Iowa | 41.692959 | -91.548043 |
| AH45 | *S. noctipennis* | Male | *H. grosseserratus* | Iowa | 43.38149 | -95.1838 |
| AH498 | *S. rugosum* | Male | *A. altissima* | Iowa | 41.665195 | -91.513398 |
| AH624 | *S. longitudinalis* | Male | *H. tuberosus* | Iowa | 41.650823 | -91.560224 |
| AH639 | *S. longitudinalis* | Male | *H. tuberosus* | Iowa | 41.650823 | -91.560224 |
| AH642 | *S. longitudinalis* | Male | *H. tuberosus* | Iowa | 41.650823 | -91.560224 |
| AH699 | *S. longitudinalis* | Male | *H. tuberosus* | Iowa | 41.692959 | -91.548043 |
| AH713 | *S. longipennis* | Male | *H. tuberosus* | Iowa | 41.651359 | -91.507785 |
| AH714 | *S. longipennis* | Male | *H. tuberosus* | Iowa | 41.651359 | -91.507785 |
| AH716 | *S. vittigera* | Male | *H. tuberosus* | Iowa | 41.651359 | -91.507785 |
| AH721 | *S. longipennis* | Male | *H. tuberosus* | Iowa | 41.651359 | -91.507785 |
| AH730 | *S. vittigera* | Male | *H. tuberosus* | Iowa | 41.651359 | -91.507785 |
| AH739 | *S. noctipennis* | Female | *H. grosseserratus* | Iowa | 41.735516 | -91.722503 |
| AH743 | *S. arculata* | Male | *H. grosseserratus* | Iowa | 41.735516 | -91.722503 |
| AH746 | *S. arculata* | Female | *H. grosseserratus* | Iowa | 41.735516 | -91.722503 |

| | | | | | | |
|---|---|---|---|---|---|---|
| AH748 | *S. noctipennis* | Female | *H. grosseserratus* | Iowa | 41.735516 | -91.722503 |
| AH75 | *"Bush's Fly"* | Male | *H. grosseserratus* | Iowa | 43.38149 | -95.1838 |
| AH751 | *S. vittigera* | Male | *H. tuberosus* | Iowa | 41.651359 | -91.507785 |
| AH751 | *S. noctipennis* | Female | *H. grosseserratus* | Iowa | 41.735516 | -91.722503 |
| AH754 | *S. longipennis* | Male | *H. tuberosus* | Iowa | 41.651359 | -91.507785 |
| AH755 | *S. longitudinalis* | Male | *H. tuberosus* | Iowa | 41.651359 | -91.507785 |
| AH756 | *S. longitudinalis* | Male | *H. tuberosus* | Iowa | 41.651359 | -91.507785 |
| AH76 | *"Bush's Fly"* | Male | *H. grosseserratus* | Iowa | 43.38149 | -95.1838 |
| AH760 | *S. longipennis* | Male | *H. tuberosus* | Iowa | 41.651359 | -91.507785 |
| AH764 | *S. longitudinalis* | Male | *H. tuberosus* | Iowa | 41.651359 | -91.507785 |
| AH767 | *S. longitudinalis* | Male | *H. tuberosus* | Iowa | 41.643284 | -91.557771 |
| AH77 | *"Bush's Fly"* | Male | *H. grosseserratus* | Iowa | 43.38149 | -95.1838 |
| AH771 | *S. longipennis* | Female | *H. tuberosus* | Iowa | 41.643284 | -91.557771 |
| AH778 | *S. longipennis* | Female | *H. tuberosus* | Iowa | 41.692959 | -91.548043 |
| AH780 | *S. longipennis* | Male | *H. tuberosus* | Iowa | 41.692959 | -91.548043 |
| AH784 | *S. longipennis* | Female | *H. tuberosus* | Iowa | 41.651359 | -91.507785 |
| AH814 | *S. longipennis* | Female | *H. tuberosus* | Iowa | 41.65288 | -91.480382 |
| AH820 | *S. longipennis* | Female | *H. tuberosus* | Iowa | 41.651359 | -91.507785 |
| AH86 | *"Bush's Fly"* | Male | *H. grosseserratus* | Iowa | 43.38149 | -95.1838 |
| AH959 | *S. longitudinalis* | Male | *H. tuberosus* | Iowa | 41.478836 | -93.910882 |
| AH971 | *S. uvedaliae* | Male | *S. uvedalia* | North Carolina | 35.92429 | -79.050225 |
| AH972 | *S. uvedaliae* | Male | *S. uvedalia* | North Carolina | 35.92429 | -79.050225 |
| AV28 | *S. intermedia* | Male | *R. laciniata* | Iowa | 43.30119 | -91.79077 |
| AY276 | *S. perfecta* | Female | *A. trifida* | Iowa | 41.65288 | -91.480382 |
| AY370 | *S. perfecta* | Male | *A. trifida* | Iowa | 41.9391 | -91.444 |
| AY533 | *S. perfecta* | Male | *A. trifida* | Iowa | 41.8863 | -91.500444 |
| AY54 | *S. intermedia* | Male | *R. laciniata* | Iowa | 43.30119 | -91.79077 |
| AY57 | *S. intermedia* | Female | *R. laciniata* | Iowa | 43.30119 | -91.79077 |
| AY88 | *S. intermedia* | Female | *R. laciniata* | Iowa | 43.30119 | -91.79077 |
| AY9 | *S. intermedia* | Female | *R. laciniata* | Iowa | 43.30119 | -91.79077 |
| AY91 | *S. intermedia* | Female | *R. laciniata* | Iowa | 43.30119 | -91.79077 |
| CAP1 | *S. longipennis* | Male | *H. tuberosus* | Maine | 43.667258 | -70.307033 |
| CAP2 | *S. longipennis* | Male | *H. tuberosus* | Maine | 43.667258 | -70.307033 |
| DH111 | *S. perfecta* | Male | *A. trifida* | Illinois | 40.113129 | -88.158744 |
| DH145 | *"Bush's Fly"* | Male | *H. grosseserratus* | Iowa | 41.01306 | -94.95483 |
| DH170 | *S. vittigera* | Female | *H. tuberosus* | Iowa | 41.421579 | -95.854282 |
| DH61 | *S. intermedia* | Male | *R. laciniata* | Iowa | 43.30119 | -91.79077 |
| DH93 | *S. longitudinalis* | Female | *H. tuberosus* | Iowa | 41.9262 | -91.4199 |
| DL21 | *S. longipennis* | Male | *H. tuberosus* | Iowa | 41.643284 | -91.557771 |
| DL9B | *S. noctipennis* | Male | *H. grosseserratus* | Iowa | 41.735516 | -91.722503 |
| EAR64 | *S. vittigera* | Male | *H. tuberosus* | Iowa | 41.65288 | -91.480382 |
| EAR82 | *S. vittigera* | Male | *H. tuberosus* | Iowa | 41.747012 | -90.515424 |
| EAR85 | *S. vittigera* | Male | *H. tuberosus* | Iowa | 41.747012 | -90.515424 |
| EAR86 | *S. vittigera* | Male | *H. tuberosus* | Iowa | 41.747012 | -90.515424 |
| ERB24 | *S. longitudinalis* | Male | *H. tuberosus* | Iowa | 41.65288 | -91.480382 |
| FR12 | *S. longipennis* | Male | *H. tuberosus* | Vermont | 43.85318 | -72.58895 |
| FR2 | *S. longipennis* | Male | *H. tuberosus* | Vermont | 43.85318 | -72.58895 |
| FR3 | *S. longipennis* | Male | *H. tuberosus* | Vermont | 43.85318 | -72.58895 |
| FW1 | *S. longipennis* | Male | *H. tuberosus* | Pennsylvania | 40.126567 | -75.170395 |
| FW2 | *S. longipennis* | Male | *H. tuberosus* | Pennsylvania | 40.126567 | -75.170395 |
| HH3 | *S. longitudinalis* | Female | *H. tuberosus* | Iowa | 41.665195 | -91.513398 |
| HHB2 | *S. longipennis* | Female | *H. tuberosus* | Iowa | 41.665195 | -91.513398 |
| HS229 | *S. noctipennis* | Female | *H. grosseserratus* | Iowa | 41.01306 | -94.95483 |
| HS70 | *S. arculata* | Male | *H. grosseserratus* | Iowa | 41.9262 | -91.4199 |
| HW44 | *S. longitudinalis* | Male | *H. tuberosus* | Iowa | 41.65288 | -91.480382 |
| JA101B | *S. longitudinalis* | Female | *H. tuberosus* | Iowa | 41.9391 | -91.444 |
| JA121 | *S. vittigera* | Female | *H. tuberosus* | Iowa | 41.421579 | -95.854282 |
| JA145 | *S. vittigera* | Female | *H. tuberosus* | Iowa | 41.421579 | -95.854282 |
| JA250 | *S. longitudinalis* | Female | *H. tuberosus* | Iowa | 41.64771 | -91.570264 |
| JA265 | *"strumosus Fly"* | Male | *H. strumosus* | Iowa | 41.915356 | -91.51616 |
| JA297 | *S. perfecta* | Male | *A. trifida* | Illinois | 40.113129 | -88.158744 |
| JA312 | *S. longitudinalis* | Female | *H. tuberosus* | Illinois | 40.113129 | -88.158744 |
| JA321 | *S. perfecta* | Male | *A. trifida* | Illinois | 40.113129 | -88.158744 |
| JA328 | *S. longitudinalis* | Female | *H. tuberosus* | Illinois | 40.113129 | -88.158744 |
| JA330 | *S. perfecta* | Female | *A. trifida* | Illinois | 40.113129 | -88.158744 |

| | | | | | | |
|---|---|---|---|---|---|---|
| JA361 | *S. vittigera* | Female | *H. tuberosus* | Illinois | 40.113129 | -88.158744 |
| JA369 | *S. perfecta* | Female | *A. trifida* | Illinois | 40.113129 | -88.158744 |
| JA47 | *S. intermedia* | Male | *R. laciniata* | Iowa | 43.30119 | -91.79077 |
| JA51 | *S. intermedia* | Male | *R. laciniata* | Iowa | 43.30119 | -91.79077 |
| JA53A | *S. intermedia* | Male | *R. laciniata* | Iowa | 43.30119 | -91.79077 |
| JA540B | *S. longipennis* | Female | *H. tuberosus* | Wisconsin | 42.88401 | -89.85717 |
| JA559 | *S. longitudinalis* | Female | *H. tuberosus* | Wisconsin | 42.88401 | -89.85717 |
| JA588B | *S. longitudinalis* | Female | *H. tuberosus* | Wisconsin | 42.88401 | -89.85717 |
| JA591B | *S. longitudinalis* | Female | *H. tuberosus* | Wisconsin | 42.88401 | -89.85717 |
| JA638B | *S. longitudinalis* | Female | *H. tuberosus* | Wisconsin | 42.88401 | -89.85717 |
| JA81 | *S. intermedia* | Male | *R. laciniata* | Illinois | 40.197027 | -88.391283 |
| JA98 | *S. vittigera* | Female | *H. tuberosus* | Iowa | 41.9391 | -91.444 |
| KNY5 | *S. longipennis* | Male | *H. tuberosus* | New York | 41.261217 | -73.687041 |
| KNY6 | *S. longipennis* | Male | *H. tuberosus* | New York | 41.261217 | -73.687041 |
| MAB3 | *S. longitudinalis* | Male | *H. tuberosus* | Iowa | 41.643284 | -91.557771 |
| MAB35 | *S. vittigera* | Male | *H. tuberosus* | Iowa | 41.651359 | -91.507785 |
| MAB36 | *S. vittigera* | Male | *H. tuberosus* | Iowa | 41.651359 | -91.507785 |
| MAB39 | *S. vittigera* | Male | *H. tuberosus* | Iowa | 41.651359 | -91.507785 |
| MAB47 | *S. longipennis* | Female | *H. tuberosus* | Iowa | 41.651359 | -91.507785 |
| MAB59 | *S. arculata* | Female | *H. grosseserratus* | Iowa | 41.735516 | -91.722503 |
| MAB62 | *S. longitudinalis* | Male | *H. tuberosus* | Iowa | 41.643284 | -91.557771 |
| MAB64A | *S. longipennis* | Female | *H. tuberosus* | Iowa | 41.651359 | -91.507785 |
| MAB65A | *S. longipennis* | Female | *H. tuberosus* | Iowa | 41.651359 | -91.507785 |
| MAB95 | *S. longipennis* | Female | *H. tuberosus* | Iowa | 41.651359 | -91.507785 |
| MCB04 | *S. longitudinalis* | Male | *H. tuberosus* | Iowa | 41.9262 | -91.4199 |
| MCB2D | *S. longipennis* | Male | *H. tuberosus* | Iowa | 41.9262 | -91.4199 |
| ME37 | *S. vittigera* | Female | *H. tuberosus* | Illinois | 40.197027 | -88.391283 |
| ME42 | *S. vittigera* | Male | *H. tuberosus* | Iowa | 41.421579 | -95.854282 |
| ME49B | *S. vittigera* | Female | *H. tuberosus* | Iowa | 41.421579 | -95.854282 |
| ME53 | *S. vittigera* | Female | *H. tuberosus* | Iowa | 41.421579 | -95.854282 |
| PG1 | *S. longipennis* | Male | *H. tuberosus* | Vermont | 41.904537 | -71.90383 |
| PG5 | *S. longipennis* | Male | *H. tuberosus* | Vermont | 41.904537 | -71.90383 |
| PP1 | *S. uvedaliae* | Male | *S. uvedalia* | Virginia | 37.550453 | -77.517475 |
| PP3 | *S. uvedaliae* | Male | *S. uvedalia* | Virginia | 37.550453 | -77.517475 |
| PP7 | *S. uvedaliae* | Female | *S. uvedalia* | Virginia | 37.550453 | -77.517475 |
| PUX3 | *S. intermedia* | Male | *R. laciniata* | Pennsylvania | 40.937221 | -78.939489 |
| RFB1 | *"Bush's Fly"* | Female | *H. grosseserratus* | Illinois | 41.913994 | -88.360007 |
| RFB6 | *"strumosus Fly"* | Female | *H. strumosus* | Illinois | 41.913994 | -88.360007 |
| SD7 | *S. longipennis* | Female | *H. tuberosus* | Iowa | 41.65288 | -91.480382 |
| SLIP12 | *S. gigantei* | Male | *H. giganteus* | Pennsylvania | 41.010882 | -80.005305 |
| SLIP13 | *S. gigantei* | Female | *H. giganteus* | Pennsylvania | 41.010882 | -80.005305 |
| SLIP14 | *S. gigantei* | Female | *H. giganteus* | Pennsylvania | 41.010882 | -80.005305 |
| SLIP15 | *S. gigantei* | Female | *H. giganteus* | Pennsylvania | 41.010882 | -80.005305 |
| SLIP19 | *S. gigantei* | Female | *H. giganteus* | Pennsylvania | 41.010882 | -80.005305 |
| SLIP4 | *S. gigantei* | Male | *H. giganteus* | Pennsylvania | 41.010882 | -80.005305 |
| SLIP5 | *S. gigantei* | Female | *H. giganteus* | Pennsylvania | 41.010882 | -80.005305 |
| SLIP6 | *S. gigantei* | Male | *H. giganteus* | Pennsylvania | 41.010882 | -80.005305 |
| SLIP7 | *S. gigantei* | Male | *H. giganteus* | Pennsylvania | 41.010882 | -80.005305 |
| SM9A | *S. perfecta* | Female | *A. trifida* | Iowa | 41.65288 | -91.480382 |
| SMAB14 | *"Bush's Fly"* | Female | *H. grosseserratus* | Illinois | 41.913994 | -88.360007 |
| SMAB19 | *"Bush's Fly"* | Female | *H. grosseserratus* | Illinois | 41.913994 | -88.360007 |
| SMAB27 | *"strumosus Fly"* | Female | *H. strumosus* | Illinois | 41.913994 | -88.360007 |
| SMAB28 | *"strumosus Fly"* | Male | *H. strumosus* | Illinois | 41.913994 | -88.360007 |
| SMAB3 | *"strumosus Fly"* | Male | *H. strumosus* | Illinois | 41.913994 | -88.360007 |
| SMAB36 | *"strumosus Fly"* | Male | *H. strumosus* | Illinois | 41.913994 | -88.360007 |
| SMAB40 | *"strumosus Fly"* | Male | *H. strumosus* | Illinois | 41.913994 | -88.360007 |
| SMAB46A | *"strumosus Fly"* | Female | *H. strumosus* | Illinois | 41.913994 | -88.360007 |
| SMAB47A | *"strumosus Fly"* | Female | *H. strumosus* | Illinois | 41.913994 | -88.360007 |
| SMAB48 | *"strumosus Fly"* | Male | *H. strumosus* | Illinois | 41.913994 | -88.360007 |
| SMAB54 | *"strumosus Fly"* | Male | *H. strumosus* | Illinois | 41.913994 | -88.360007 |
| SMAB55 | *"strumosus Fly"* | Male | *H. strumosus* | Illinois | 41.913994 | -88.360007 |
| st0015 | *S. longitudinalis* | Female | *H. tuberosus* | Iowa | 41.9654 | -91.5874 |
| st0019 | *S. longitudinalis* | Male | *H. tuberosus* | Iowa | 41.9654 | -91.5874 |
| st0024 | *S. longitudinalis* | Female | *H. tuberosus* | Iowa | 41.9654 | -91.5874 |
| st0028 | *S. vittigera* | Female | *H. tuberosus* | Iowa | 41.9654 | -91.5874 |

| | | | | | | |
|---|---|---|---|---|---|---|
| st0065 | *S. longitudinalis* | Female | *H. tuberosus* | Iowa | 41.9654 | -91.5874 |
| st0072 | *S. longitudinalis* | Female | *H. tuberosus* | Iowa | 41.9654 | -91.5874 |
| st0087 | *S. longitudinalis* | Male | *H. tuberosus* | Iowa | 41.9654 | -91.5874 |
| st0091 | *S. longipennis* | Male | *H. tuberosus* | Iowa | 41.9654 | -91.5874 |
| st0093 | *S. longitudinalis* | Male | *H. tuberosus* | Iowa | 41.9654 | -91.5874 |
| st0100 | *S. longipennis* | Male | *H. tuberosus* | Iowa | 41.9654 | -91.5874 |
| st0101 | *S. longipennis* | Male | *H. tuberosus* | Iowa | 41.9654 | -91.5874 |
| st0104 | *S. longipennis* | Male | *H. tuberosus* | Iowa | 41.9654 | -91.5874 |
| st0105 | *S. longipennis* | Male | *H. tuberosus* | Iowa | 41.9654 | -91.5874 |
| st0106 | *S. longipennis* | Male | *H. tuberosus* | Iowa | 41.9654 | -91.5874 |
| st0108 | *S. longipennis* | Male | *H. tuberosus* | Iowa | 41.9654 | -91.5874 |
| st0111 | *S. longipennis* | Female | *H. tuberosus* | Iowa | 41.9654 | -91.5874 |
| st0112 | *S. longipennis* | Female | *H. tuberosus* | Iowa | 41.9654 | -91.5874 |
| st0113 | *S. longitudinalis* | Male | *H. tuberosus* | Iowa | 41.9654 | -91.5874 |
| st0114 | *S. longitudinalis* | Male | *H. tuberosus* | Iowa | 41.9262 | -91.4199 |
| st0115 | *S. longitudinalis* | Male | *H. tuberosus* | Iowa | 41.65288 | -91.480382 |
| st0116 | *S. longitudinalis* | Male | *H. tuberosus* | Iowa | 41.65288 | -91.480382 |
| st0118 | *S. longipennis* | Male | *H. tuberosus* | Iowa | 41.65288 | -91.480382 |
| st0130 | *S. longitudinalis* | Male | *H. tuberosus* | Iowa | 41.65288 | -91.480382 |
| st0132 | *S. longitudinalis* | Male | *H. tuberosus* | Iowa | 41.65288 | -91.480382 |
| st0134 | *S. longitudinalis* | Male | *H. tuberosus* | Iowa | 41.65288 | -91.480382 |
| st0135 | *S. longitudinalis* | Female | *H. tuberosus* | Iowa | 41.9262 | -91.4199 |
| st0136 | *S. longipennis* | Male | *H. tuberosus* | Iowa | 41.9654 | -91.5874 |
| st0137 | *S. longipennis* | Female | *H. tuberosus* | Iowa | 41.9654 | -91.5874 |
| st0139 | *S. longipennis* | Male | *H. tuberosus* | Iowa | 41.9654 | -91.5874 |
| TC1 | *S. noctipennis* | Male | *H. grosseserratus* | Iowa | 41.747012 | -90.515424 |
| TC104 | *S. vittigera* | Female | *H. tuberosus* | Iowa | 41.01306 | -94.95483 |
| TC14 | *S. arculata* | Male | *H. grosseserratus* | Iowa | 41.747012 | -90.515424 |
| TC146 | *S. vittigera* | Female | *H. tuberosus* | Iowa | 41.9654 | -91.5874 |
| TC15 | *S. arculata* | Male | *H. grosseserratus* | Iowa | 41.747012 | -90.515424 |
| TC151 | *S. longitudinalis* | Female | *H. tuberosus* | Iowa | 41.9262 | -91.4199 |
| TC156B | *S. longitudinalis* | Female | *H. tuberosus* | Iowa | 41.9262 | -91.4199 |
| TC168 | *S. vittigera* | Female | *H. tuberosus* | Iowa | 41.9262 | -91.4199 |
| TC16A | *S. longitudinalis* | Female | *H. tuberosus* | Iowa | 41.9262 | -91.4199 |
| TC176 | *S. longitudinalis* | Female | *H. tuberosus* | Iowa | 41.9654 | -91.5874 |
| TC2 | *S. noctipennis* | Male | *H. grosseserratus* | Iowa | 41.747012 | -90.515424 |
| TC232 | *S. perfecta* | Female | *A. trifida* | Illinois | 40.113129 | -88.158744 |
| TC254 | *S. vittigera* | Female | *H. tuberosus* | Illinois | 40.113129 | -88.158744 |
| tc256 | *S. perfecta* | Male | *A. trifida* | Illinois | 40.113129 | -88.158744 |
| TC262A | *S. perfecta* | Male | *A. trifida* | Illinois | 40.113129 | -88.158744 |
| TC266 | *S. perfecta* | Male | *A. trifida* | Illinois | 40.113129 | -88.158744 |
| TC276 | *S. longitudinalis* | Female | *H. tuberosus* | Illinois | 40.113129 | -88.158744 |
| TC3 | *S. noctipennis* | Male | *H. grosseserratus* | Iowa | 41.747012 | -90.515424 |
| TC340 | *"Bush's Fly"* | Male | *H. grosseserratus* | Iowa | 41.01306 | -94.95483 |
| TC4 | *S. noctipennis* | Male | *H. grosseserratus* | Iowa | 41.747012 | -90.515424 |
| TC415 | *S. longitudinalis* | Female | *H. tuberosus* | Wisconsin | 42.88401 | -89.85717 |
| TC437B | *S. longitudinalis* | Female | *H. tuberosus* | Wisconsin | 42.88401 | -89.85717 |
| TC54 | *S. intermedia* | Male | *R. laciniata* | Iowa | 43.30119 | -91.79077 |
| TC55 | *S. intermedia* | Male | *R. laciniata* | Iowa | 43.30119 | -91.79077 |
| TC9 | *S. noctipennis* | Male | *H. grosseserratus* | Iowa | 41.747012 | -90.515424 |
| TC91B | *S. intermedia* | Male | *R. laciniata* | Illinois | 40.197027 | -88.391283 |
| VT3 | *S. longipennis* | Male | *H. tuberosus* | Vermont | 44.256938 | -72.517137 |
| WC29 | *S. longitudinalis* | Female | *H. tuberosus* | Iowa | 41.64771 | -91.570264 |
| WC78 | *S. longitudinalis* | Female | *H. tuberosus* | Iowa | 41.64771 | -91.570264 |
| WCP05 | *S. longitudinalis* | Male | *H. tuberosus* | Iowa | 41.64771 | -91.570264 |
| WCP130019 | *S. vittigera* | Female | *H. tuberosus* | Iowa | 41.64771 | -91.570264 |
| WCP13006 | *S. vittigera* | Female | *H. tuberosus* | Iowa | 41.64771 | -91.570264 |
| WCP2 | *S. longitudinalis* | Female | *H. tuberosus* | Iowa | 41.64771 | -91.570264 |
| WCP28 | *S. longipennis* | Female | *H. tuberosus* | Iowa | 41.64771 | -91.570264 |
| WCP29 | *S. longipennis* | Female | *H. tuberosus* | Iowa | 41.64771 | -91.570264 |
| WCP7 | *S. longipennis* | Female | *H. tuberosus* | Iowa | 41.64771 | -91.570264 |

Supplemental Table 2. Pairwise t-tests comparing male and female wing centroid size. P-values in bold indicate significant size differences between males and females of the same species following Bonferroni correction.

| **Species (Male vs Female)** | P-Value | Males (N) | Females (N) |
|---|---|---|---|
| *S. intermedia* | 0.40239783 | 11 | 5 |
| *S. gigantei* | 0.935260694 | 4 | 5 |
| *S. longitudinalis* | **0.004070414** | 27 | 25 |
| *S. longipennis* | 0.441285724 | 35 | 18 |
| *S. arculata* | **0.000503267** | 15 | 4 |
| *S. vittigera* (*H. tuberosus*) | 0.038579489 | 9 | 15 |
| *"strumosus Fly"* | 0.230960761 | 8 | 4 |
| "Bush's Fly" | **0.009975878** | 16 | 4 |
| *S. noctipennis* | 0.142923889 | 8 | 6 |
| *S. uvedaliae* | 0.087098221 | 5 | 2 |
| *S. perfecta* | **0.011230084** | 9 | 5 |

Supplemental Table 3. Table reporting the p-values of all pairwise t-tests comparing the centroid size of *Strauzia* wings. The top right section of the table reports all possible comparisons among *Strauzia* females and the bottom left reports all possible comparisons among *Strauzia* males. Values in bold are significantly different following a Bonferroni correction.

Female Pairwise Comparisons

Male Pairwise Comparisons

| | *S. arculata* | *"Bush' Fly"* | *S. gigantei* | *S. intermedia* | *S. longipennis* | *S. longitudinalis* | *S. noctipennis* | *S. perfecta* | "*strumosus* fly" | *S. uvedaliae* | *S. vittigera* |
|---|---|---|---|---|---|---|---|---|---|---|---|
| *S. arculata* | | 0.043988849 | 0.230155041 | 0.0037186 | 0.13543422 | 0.008206789 | 0.003455285 | 0.681716247 | 0.880153437 | 0.467034672 | 0.213600491 |
| *"Bush's Fly"* | 0.280307966 | | 0.706842658 | **0.000810375** | 0.104188323 | 0.254055943 | 0.708940053 | 0.03062146 | 0.181071926 | 0.929175974 | 0.092676404 |
| *S. gigantei* | **6.44158E-05** | **0.000119147** | | 0.011047436 | 0.44873995 | 0.719365831 | 0.524448291 | 0.103618641 | 0.388076153 | 0.641557145 | 0.418797107 |
| *S. intermedia* | 0.018973046 | 0.002951605 | **6.24017E-06** | | **0.000596792** | **0.00024219** | **5.476E-05** | 0.003563993 | 0.048958155 | 0.000475493 | **0.000631013** |
| *S. longipennis* | **1.03334E-05** | 0.002923334 | 0.009824615 | **2.35295E-07** | | 0.302686983 | 0.013985807 | 0.133278683 | 0.709017414 | 0.024632553 | 0.886608228 |
| *S. longitudinalis* | **0.000790756** | 0.049808502 | 0.002173332 | **3.51511E-06** | 0.158799 | | 0.06066585 | 0.01866515 | 0.436286609 | 0.084775746 | 0.25871803 |
| *S. noctipennis* | **3.17392E-05** | **0.000214822** | 0.600081088 | **1.40246E-06** | 0.026135589 | 0.004472786 | | 0.002369258 | 0.120315029 | 0.684902665 | 0.012179142 |
| *S. perfecta* | 0.694938267 | 0.165090963 | **6.10825E-05** | 0.048613903 | **2.78274E-05** | **0.000625262** | **2.62326E-05** | | 0.773263739 | 0.010026814 | 0.185865312 |
| "*strumosus* fly" | 0.953649431 | 0.460752153 | **0.000695103** | 0.161531751 | 0.01688635 | 0.057181988 | 0.001158332 | 0.861593331 | | 0.768763015 | 0.75648731 |
| *S. uvedaliae* | 0.04035366 | 0.091718395 | 0.190176706 | 0.008300355 | 0.732887532 | 0.378620461 | 0.314831842 | 0.031565121 | 0.048863342 | | 0.020389524 |
| *S. vittigera* | 0.639562084 | 0.843300655 | 0.002025064 | 0.077894309 | 0.055460925 | 0.155993856 | 0.004048937 | 0.497830235 | 0.673938978 | 0.100930448 | |

Supplemental Table 4. Table reporting the p-values of all pairwise t-tests comparing the centroid size of *Strauzia* male wings after scaling wing size by the length of the average forefemur length of each species. Values in bold are significantly different following a Bonferroni correction.

| | *S. arculata* | *"Bush's Fly"* | *S. intermedia* | *S. longipennis* | *S. longitudinalis* | *S. noctipennis* | *S. perfecta* | *S. vittigera* |
|---|---|---|---|---|---|---|---|---|
| *S. arculata* | | | | | | | | |
| *"Bush's Fly"* | 0.91355211 | | | | | | | |
| *S. intermedia* | 0.429234862 | 0.41132815 | | | | | | |
| *S. longipennis* | 0.067285521 | 0.088416523 | 0.471923061 | | | | | |
| *S. longitudinalis* | 0.002084029 | 0.004579877 | 0.050439713 | 0.070033199 | | | | |
| *S. noctipennis* | **9.30915E-05** | **8.69786E-05** | **0.000549837** | **0.000989954** | 0.010068718 | | | |
| *S. perfecta* | 0.505877347 | 0.480981263 | 0.85925358 | 0.309581553 | 0.021374522 | **0.000343965** | | |
| *S. vittigera* | 0.309002809 | 0.294864628 | 0.588735348 | 0.860925066 | 0.534291064 | 0.032013296 | 0.508996014 | |

Supplemental Table 5. P-values from MANOVAs comparing the wing shapes of all males or all females that share the same host plant species. P-values in bold indicate those that are significant following a Bonferroni correction.

*H. grosseserratus* Males

| | *S. arculata* | "Bush's Fly" |
|---|---|---|
| *S. arculata* | | |
| "Bush's Fly" | **2.3564E-09** | |
| *S. noctipennis* | **7.1834E-11** | **1.5471E-08** |

*H. grosseserratus* Females

| | *S. arculata* | "Bush's Fly" |
|---|---|---|
| *S. arculata* | | |
| "Bush's Fly" | **0.010598** | |
| *S. noctipennis* | **0.0040119** | 0.48127 |

*H. tuberosus* Males

| | *S. longipennis* | *S. longitudinalis* |
|---|---|---|
| *S. longipennis* | | |
| *S. longitudinalis* | **5.5431E-07** | |
| *S. vittigera* | **9.4817E-08** | **1.2014E-06** |

*H. tuberosus* Females

| | *S. longipennis* | *S. longitudinalis* |
|---|---|---|
| *S. longipennis* | | |
| *S. longitudinalis* | **0.0027827** | |
| *S. vittigera* | 0.027044 | 0.027811 |

Supplemental Table 6. P-values of a MANOVA comparison of the wing shape of all male *Strauzia* species included in the analysis. Values in bold indicate significant shape differences following a Bonferroni correction.

| | *S. arculata* | *"Bush's Fly"* | *S. gigantei* | *S. intermedia* | *S. longipennis* | *S. longitudinalis* | *S. noctipennis* | *S. perfecta* | *"strumosus Fly"* | *S. uvedaliae* | *S. verbesinae* |
|---|---|---|---|---|---|---|---|---|---|---|---|
| *S. arculata* | | | | | | | | | | | |
| "Bush's Fly" | **1.24E-07** | | | | | | | | | | |
| *S. gigantei* | 0.0021126 | **4.52E-07** | | | | | | | | | |
| *S. intermedia* | **3.06E-08** | **4.93E-13** | 0.047889 | | | | | | | | |
| *S. longipennis* | **1.67E-07** | **4.92E-13** | **7.93E-07** | **9.86E-18** | | | | | | | |
| *S. longitudinalis* | **1.07E-09** | **8.73E-14** | **3.81E-06** | **2.42E-15** | **0.00012644** | | | | | | |
| *S. noctipennis* | **4.60E-09** | **4.75E-08** | **3.60E-05** | **1.15E-10** | **4.18E-10** | **3.15E-10** | | | | | |
| *S. perfecta* | **7.01E-07** | **1.35E-10** | 0.018396 | **4.61E-07** | **1.27E-07** | **6.34E-05** | **4.37E-07** | | | | |
| *"strumosus Fly"* | 0.56102 | **2.99E-06** | 0.037297 | **9.65E-06** | **0.00065221** | **1.18E-05** | **4.52E-06** | **0.0004497** | | | |
| *S. uvedaliae* | 0.15979 | **9.30E-05** | 0.069375 | **4.77E-05** | 0.22566 | 0.012656 | **0.00032961** | 0.0096224 | 0.6422 | | |
| *S. verbesinae* | 0.0020764 | **1.18E-05** | 0.38314 | 0.72793 | **1.02E-07** | **5.42E-07** | **0.00030031** | 0.0033762 | 0.021447 | 0.075054 | |
| *S. vittigera* | **3.02E-05** | **2.73E-10** | 0.25504 | **1.89E-05** | **5.16E-08** | **2.85E-07** | **1.94E-07** | 0.035857 | 0.0063305 | 0.02514 | 0.013164 |

Supplemental Table 7. P-values of a MANOVA comparison of the wing shape of all female *Strauzia* species included in the analysis. Values in bold indicate significant shape differences following a Bonferroni correction.

| | *S. arculata* | "*Bush's Fly*" | *S. gigantei* | *S. intermedia* | *S. longipennis* | *S. longitudinalis* | *S. noctipennis* | *S. perfecta* | "*strumosus* fly" | *S. uvedaliae* |
|---|---|---|---|---|---|---|---|---|---|---|
| *S. arculata* | | | | | | | | | | |
| "*Bush's Fly*" | 0.13731 | | | | | | | | | |
| *S. gigantei* | 0.047646 | 0.064846 | | | | | | | | |
| *S. intermedia* | 0.029815 | 0.023393 | 0.49647 | | | | | | | |
| *S. longipennis* | **3.81E-05** | 0.014792 | 0.0033764 | **1.68E-05** | | | | | | |
| *S. longitudinalis* | **4.30E-08** | 0.0020608 | **9.29E-05** | **1.39E-07** | 0.0034954 | | | | | |
| *S. noctipennis* | 0.056594 | 0.77693 | 0.0083435 | 0.0022066 | **0.0001496** | **1.59E-06** | | | | |
| *S. perfecta* | 0.040716 | 0.097968 | 0.22795 | 0.050017 | 0.37299 | 0.014897 | 0.012546 | | | |
| "*strumosus* fly" | 0.19795 | 0.25046 | 0.23595 | 0.08878 | 0.018672 | **0.00012634** | 0.094429 | 0.21088 | | |
| *S. uvedaliae* | 0.60481 | 0.68872 | 0.89699 | 0.54659 | 0.28254 | 0.036631 | 0.2753 | 0.70191 | 0.89924 | |
| *S. vittigera* | **5.71E-05** | 0.058673 | 0.0089477 | **9.28E-05** | 0.014396 | 0.014509 | 0.00099276 | 0.020715 | 0.01838 | 0.36268 |

**Supplemental Figure 1.** Warp grids showing the difference in shape from the average female wing to the average male wing for each *Strauzia* species. Numbered points correspond to the wing landmarks used throughout the study (Figure 2).

**Supplemental Figure 2.** Principal component analysis of all male Strauzia wings included in this study. Axes represent PC1 and PC2. Points, lines encompassing points, and species labels are color coded to indicate which points are associated with each species.

**Supplemental Figure 3.** Principal component analysis of all female Strauzia wings included in this study. Axes represent PC1 and PC2. Points, lines encompassing points, and species labels are color coded to indicate which points are associated with each species.

**Supplemental Figure 4.** Principal component analysis of male *Strauzia* wings that share the plant host, *H. tuberosus*, including *S. longipennis*, *S. vittigera*, and *S. longitudinalis*. Axes represent PC1 and PC2. Points, lines encompassing points, and species labels are color coded to indicate which points are associated with each species.

**Supplemental Figure 5.** Principal component analysis of male *Strauzia* wings that share the plant host, *H. grosseserratus*, including *S. arculata*, *S. noctipennis*, and "Bush's Fly". Axes represent PC1 and PC2. Points, lines encompassing points, and species labels are color coded to indicate which points are associated with each species.

**Supplemental Figure 6.** Principal component analysis of female *Strauzia* wings that share the plant host, *H. grosseserratus*, including *S. arculata*, *S. noctipennis*, and "Bush's Fly". Axes represent PC1 and PC2. Points, lines encompassing points, and species labels are color coded to indicate which points are associated with each species.

**Supplemental Figure 7.** Principal component analysis of female *Strauzia* wings that share the plant host, *H. tuberosus*, including *S. longipennis*, *S. vittigera*, and *S. longitudinalis*. Axes represent PC1 and PC2. Points, lines encompassing points, and species labels are color coded to indicate which points are associated with each species.

Supplemental Figure 1

S. arculata

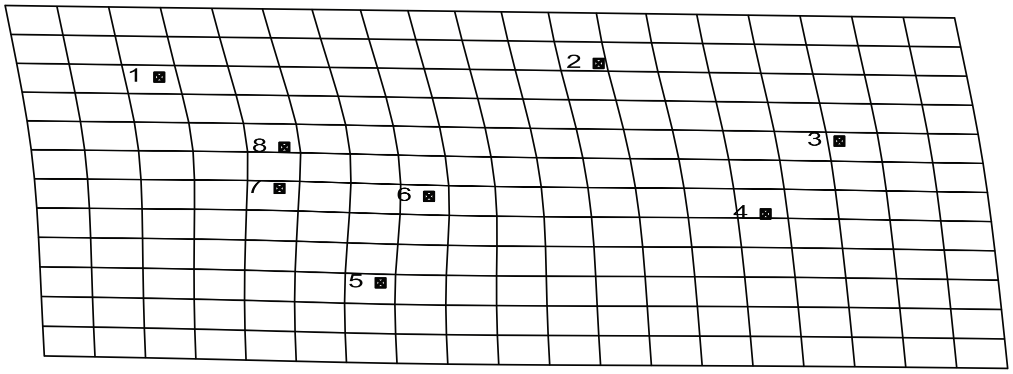


S. intermedia

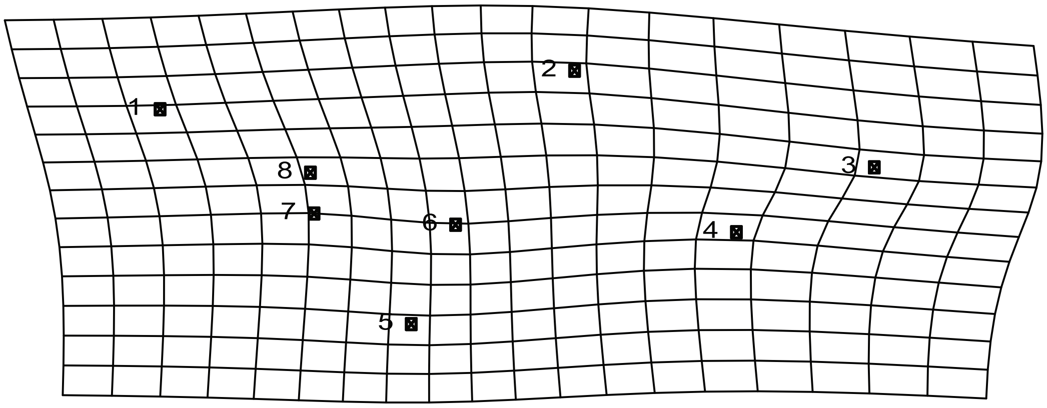


S. perfecta

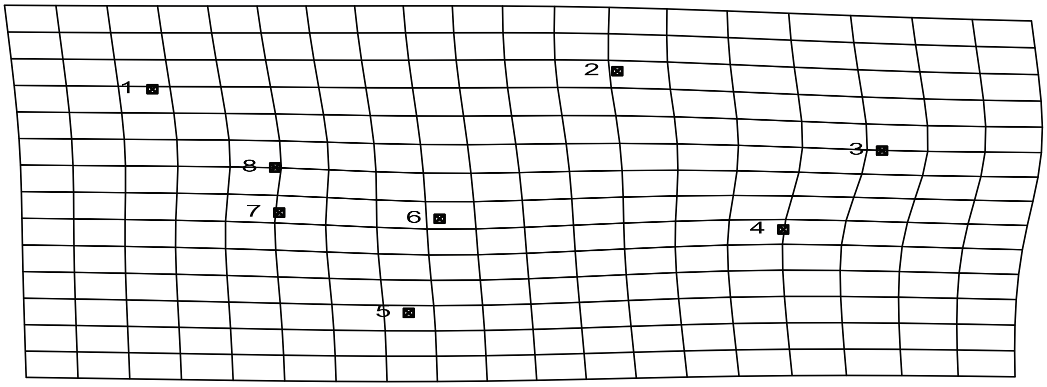


S. vittigera

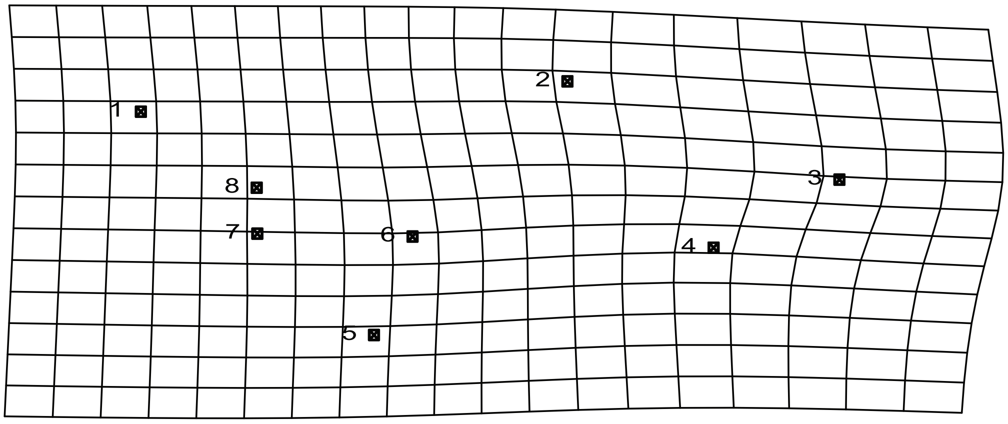

"strumosus Fly"

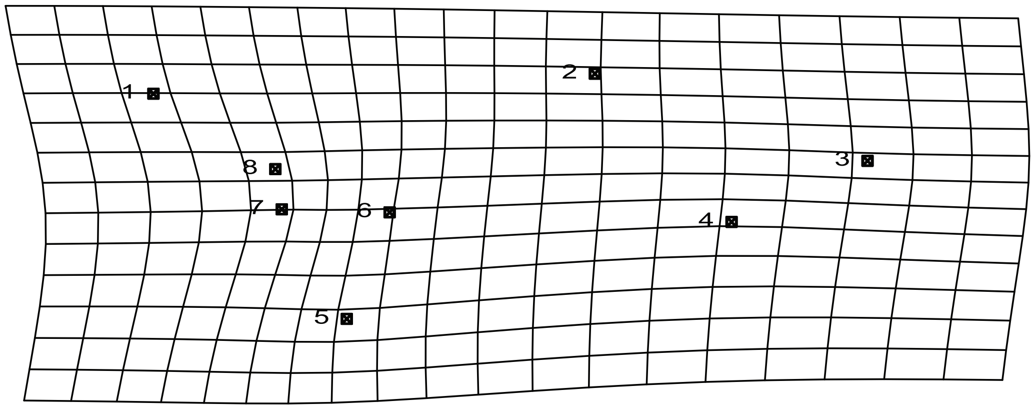


S. noctipennis

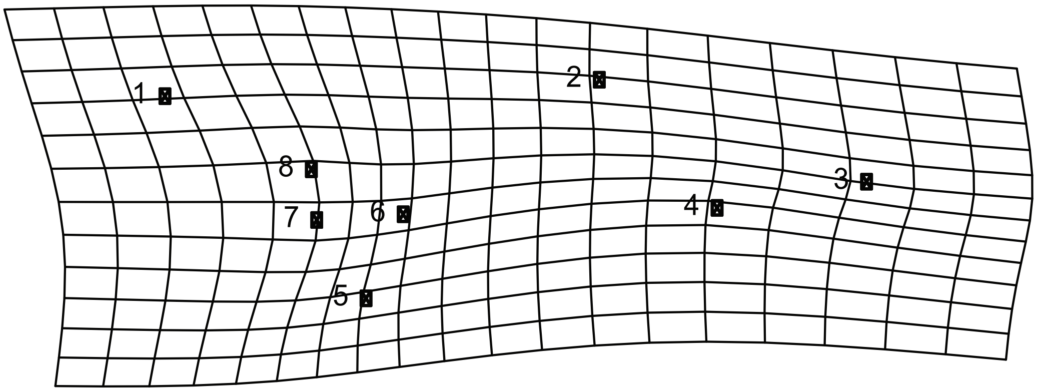


S. longitudinalis

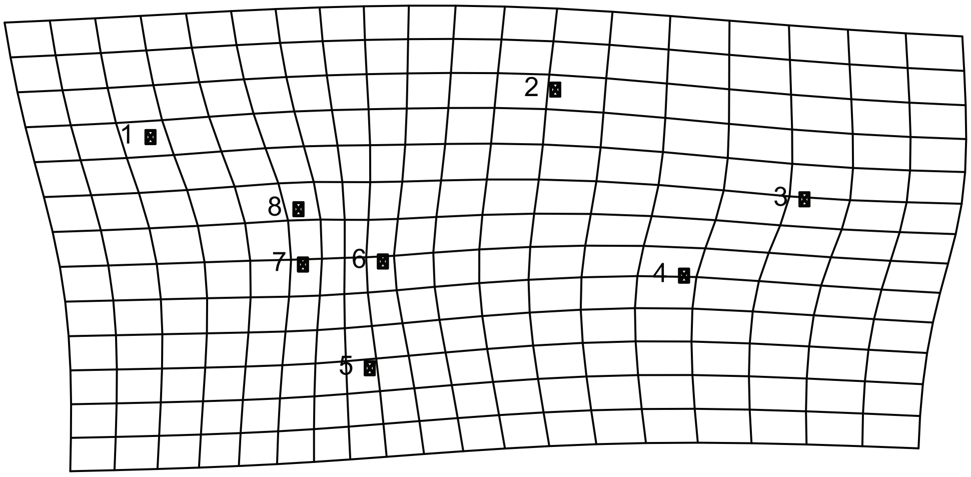

S. longipennis

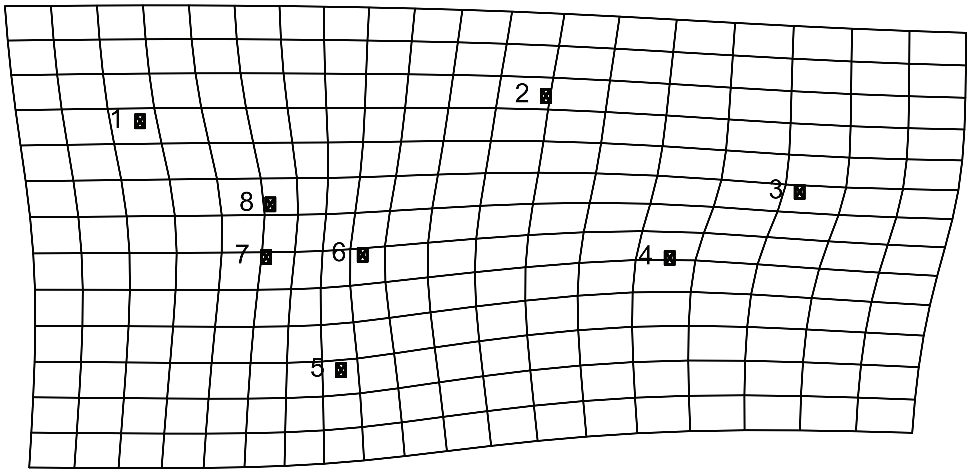


S. gigantei

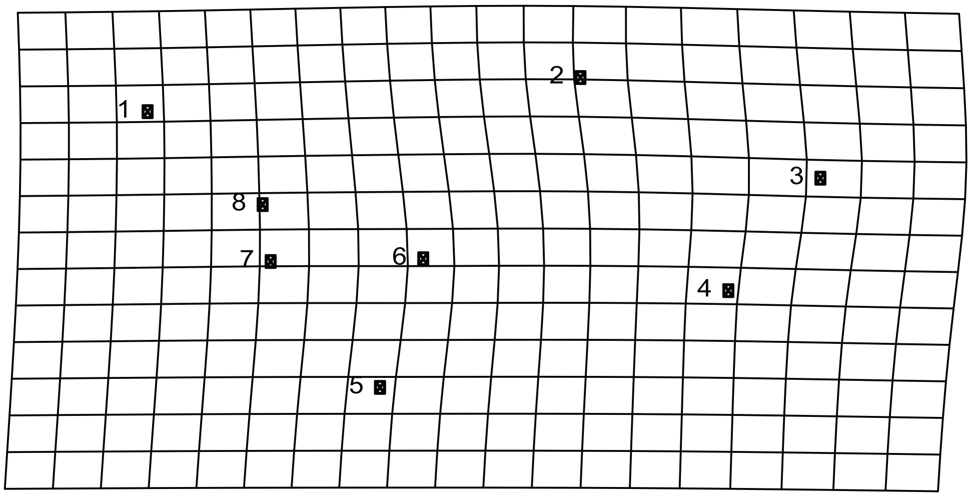


“Bush’s Fly”

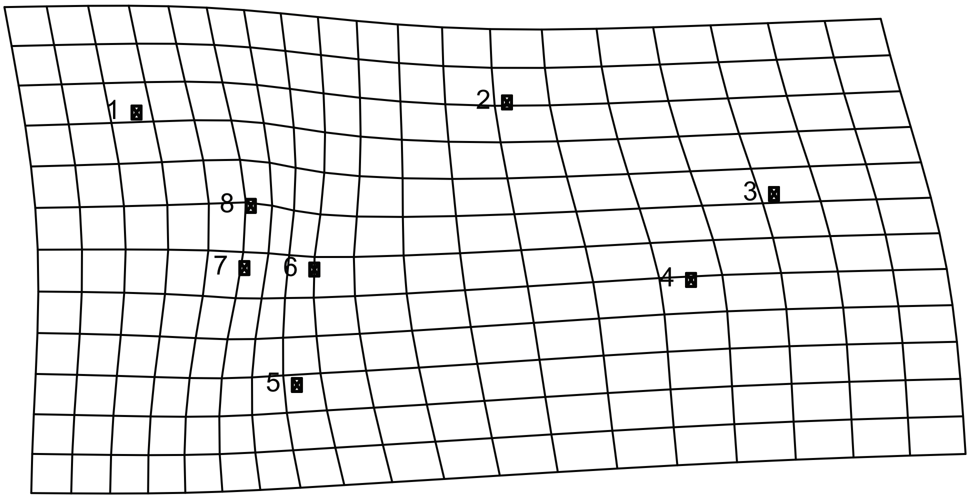

Supplemental Figure 2

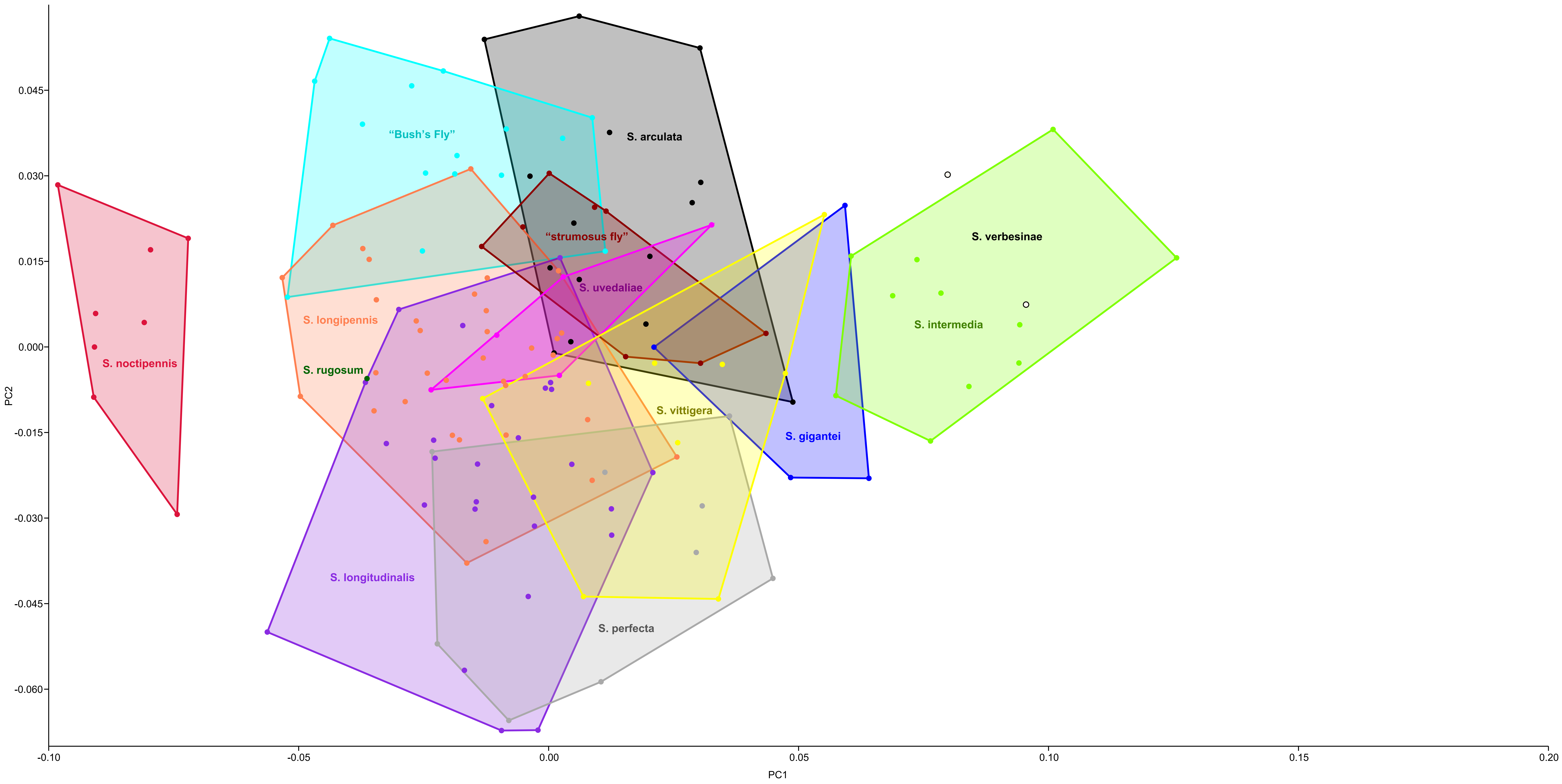

Supplemental Figure 3

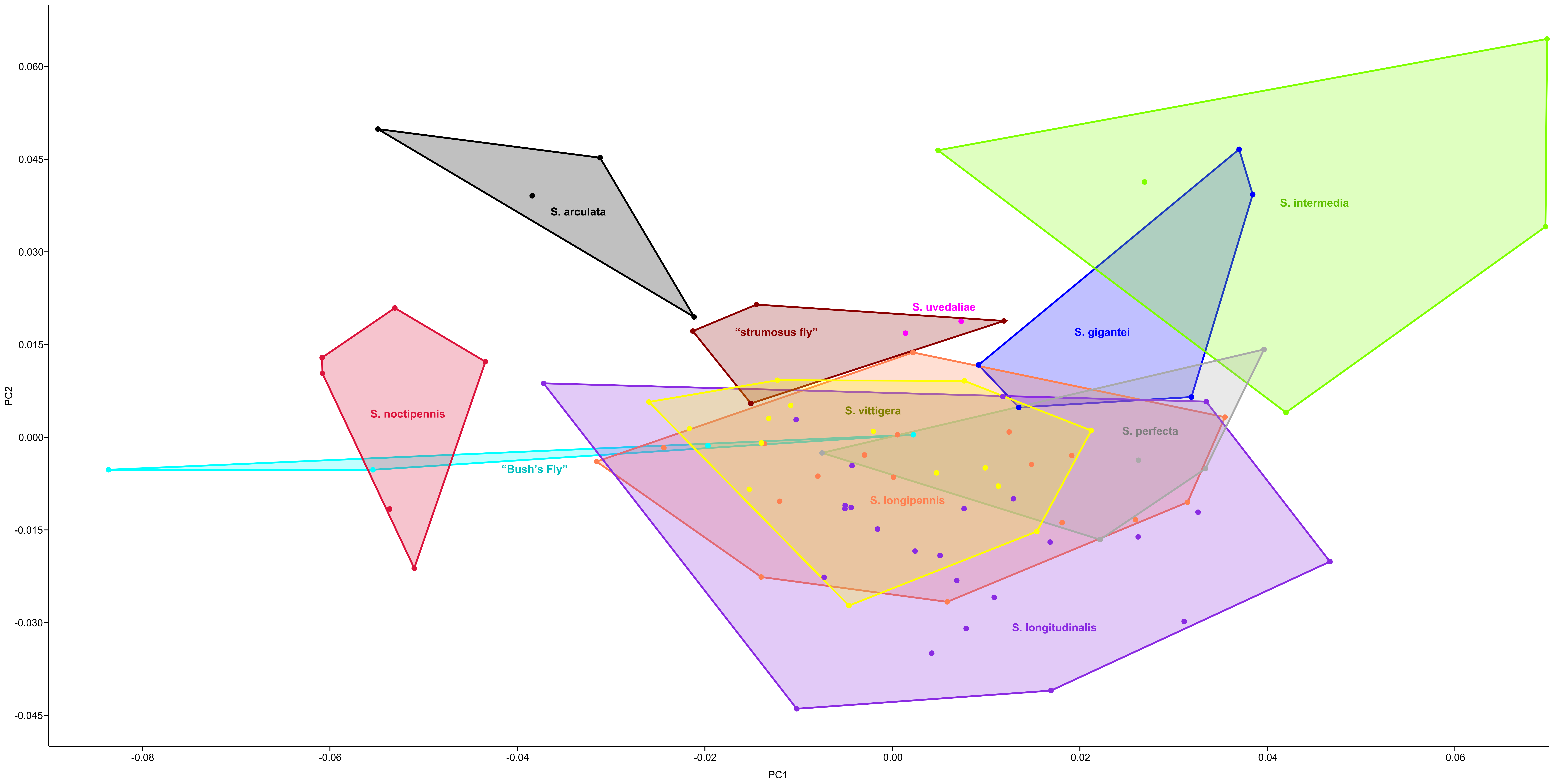

Supplemental Figure 4

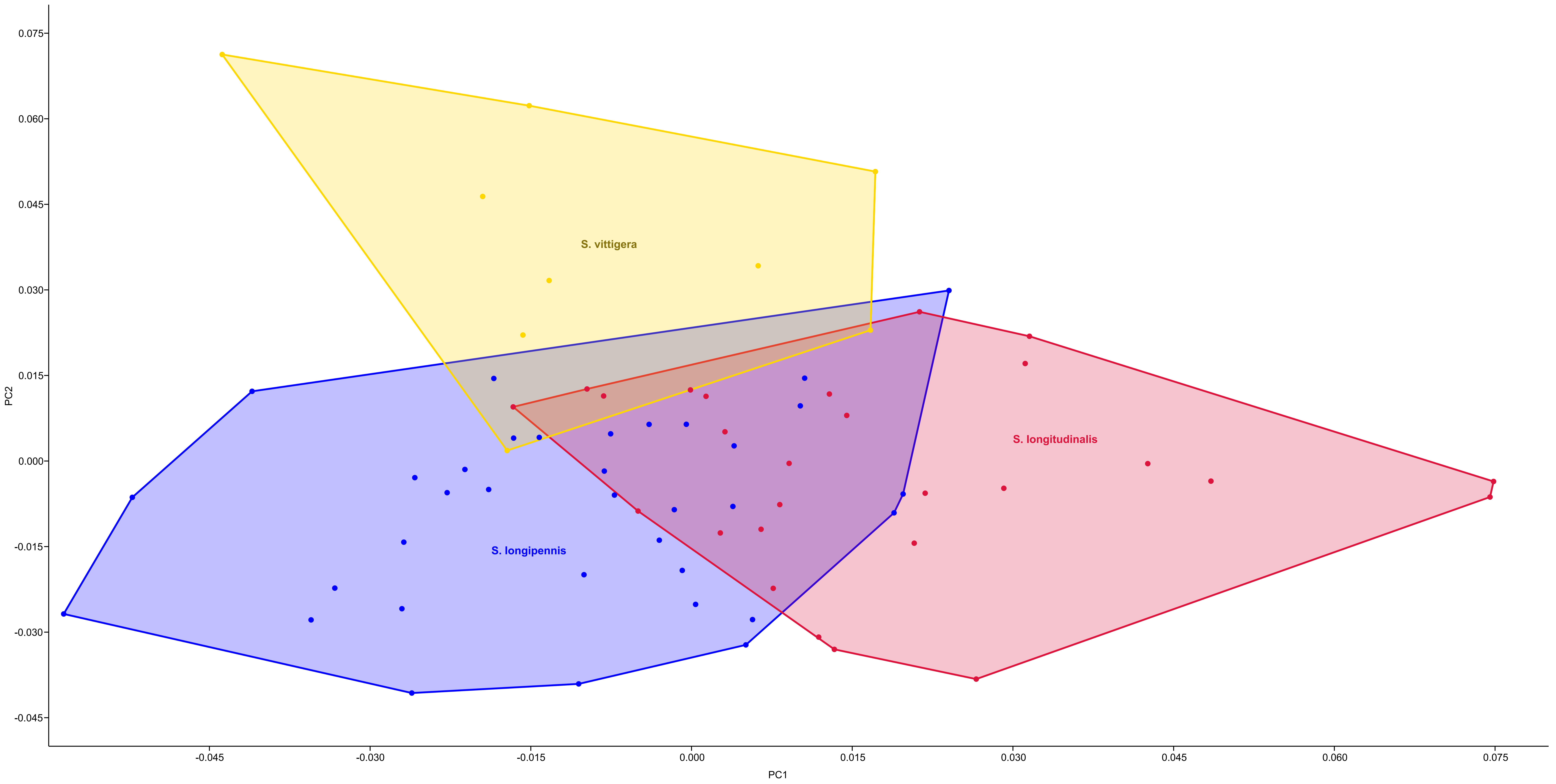

Supplemental Figure 5

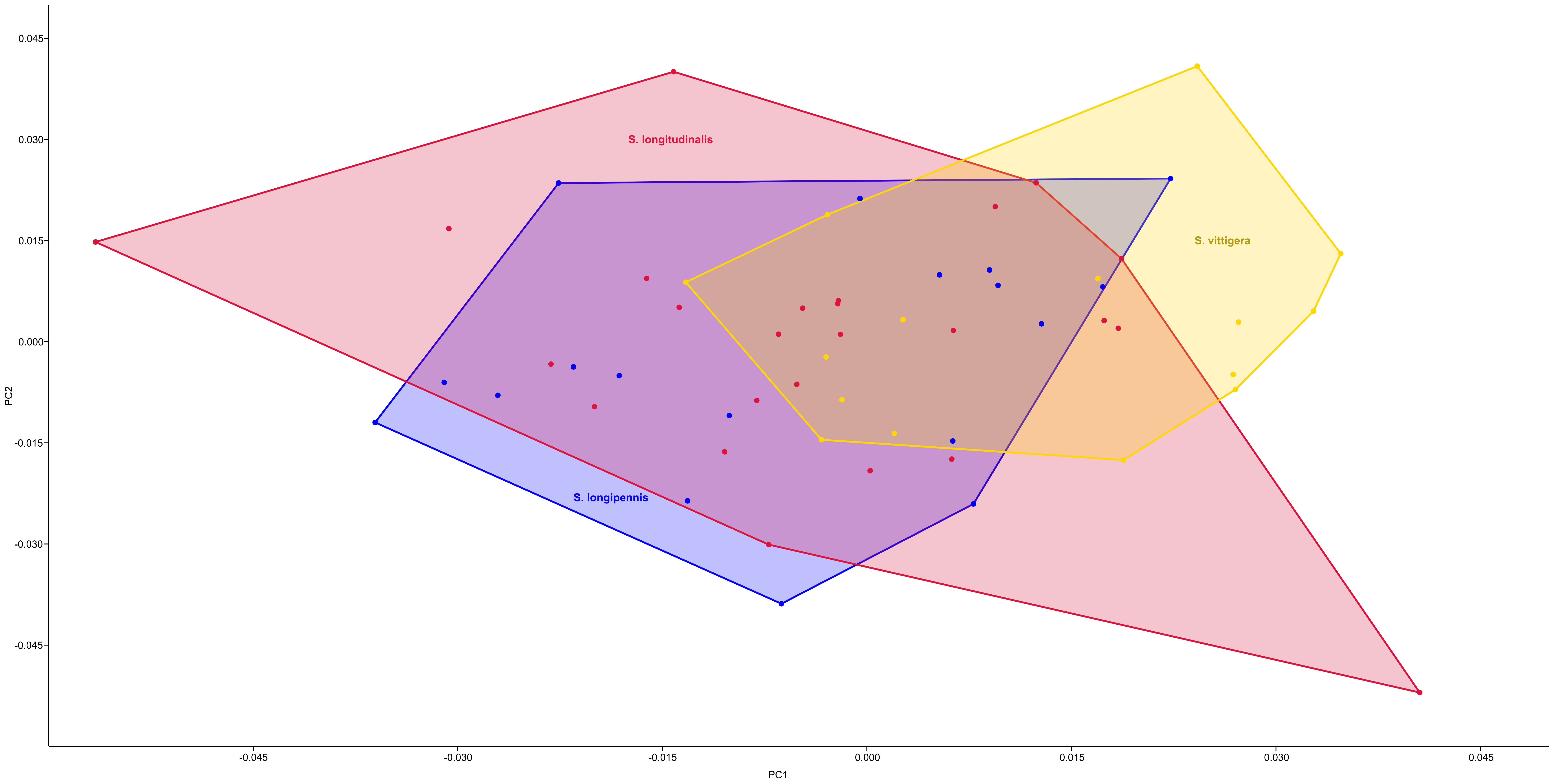

Supplemental Figure 6

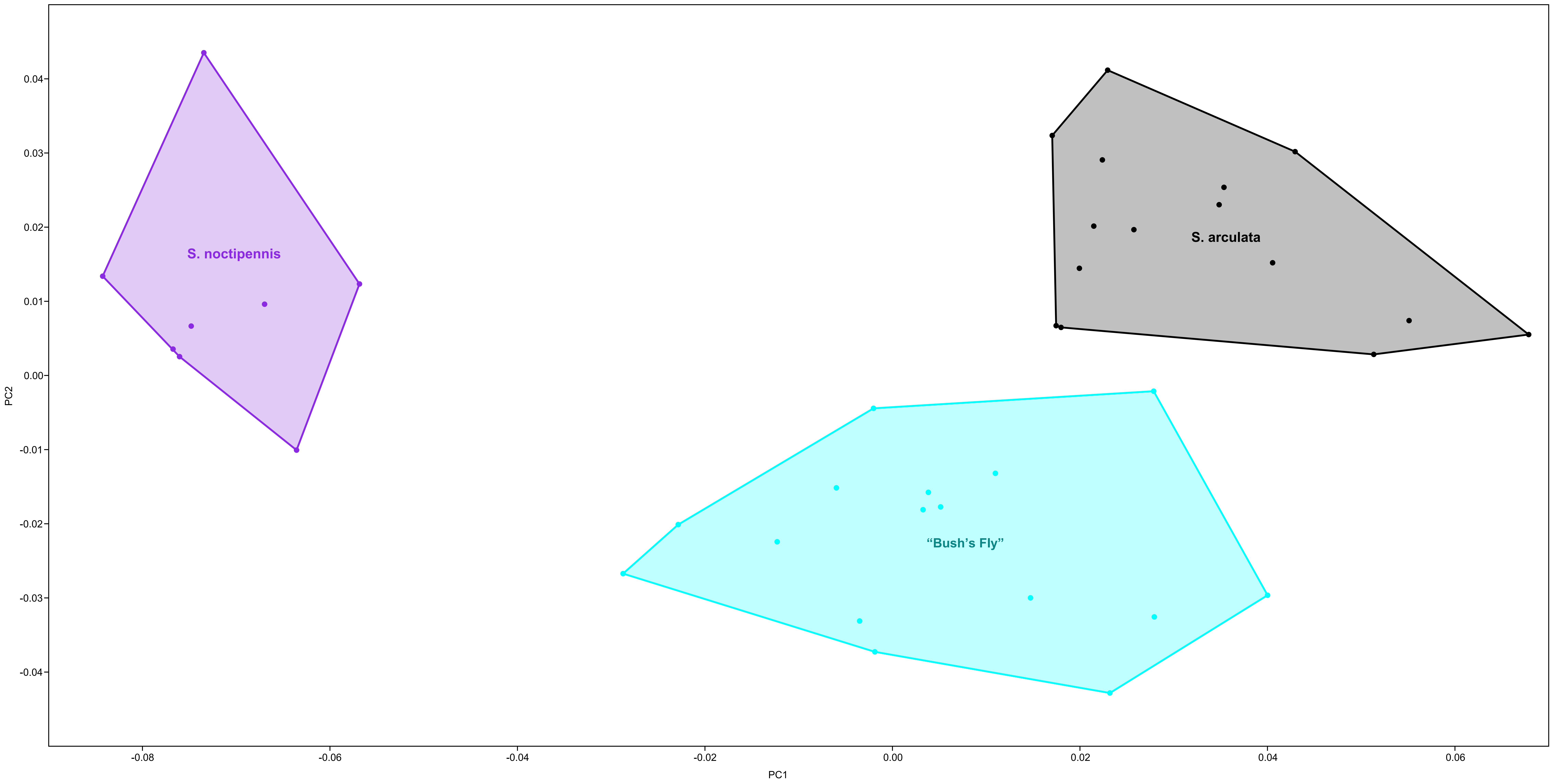

Supplemental Figure 7

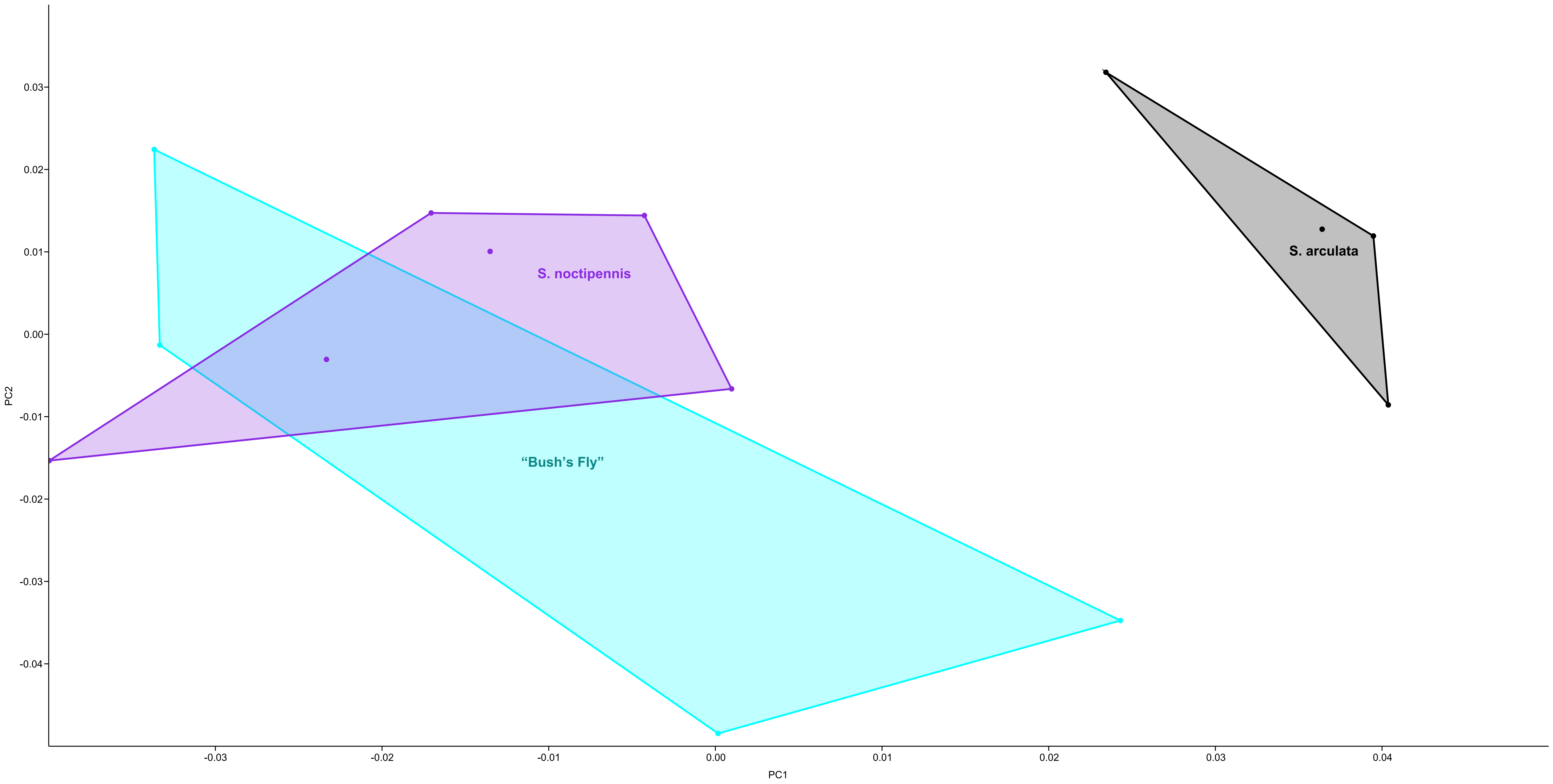
